# Biomechanical property of limbal niche maintains stemness through YAP

**DOI:** 10.1101/2021.05.25.445490

**Authors:** Swarnabh Bhattacharya, Abhishek Mukherjee, Sabrina Pisano, Shalini Dimri, Eman Knaane, Anna Altshuler, Waseem Nasser, Sunanda Dey, Lidan Shi, Ido Mizrahi, Ophir Jokel, Aya Amitai-Lange, Anna Kaganovsky, Michael Mimouni, Sergiu Socea, Peleg Hasson, Chloe Feral, Haguy Wolfenson, Ruby Shalom-Feuerstein

## Abstract

Stem cells’ (SCs) decision to self-renew or differentiate largely depends on the external control of their niche. However, the complex mechanisms that underlie this crosstalk are poorly understood. To address this question, we focused on the corneal epithelial SC model in which the SC niche, known as the limbus, is spatially segregated from the differentiation compartment. We report that the unique biomechanical property of the limbus supports the nuclear localization and function of Yes-associated protein (YAP), a putative mediator of the mechanotransduction pathway. Perturbation of tissue stiffness or YAP activity affects SC function as well as tissue integrity under homeostasis and significantly inhibited the regeneration of the SC population following SC depletion. *In vitro* experiments revealed that substrates with the rigidity of the corneal differentiation compartment inhibit YAP localization and induce differentiation, a mechanism that is mediated by the TGFβ−SMAD2/3 pathway. Taken together, these results indicate that SC sense biomechanical niche signals and that manipulation of mechano-sensory machinery or its downstream biochemical output may bear fruits in SC expansion for regenerative therapy.

**Highlights:**

- YAP is essential for limbal SC function, regeneration, and dedifferentiation
- Lox over-expression stiffens the limbal niche, affects SC phenotype and corneal integrity
- Corneal rigidity represses YAP and stemness in a SMAD2/3-dependent manner
- Manipulation of mechanosensory or TGF-β pathway influences limbal SC expansion *in vitro*

## Introduction

Stem cell (SC) fate decisions are controlled by extrinsic signals derived from the local microenvironment, known as the niche (Spradling *et al*, 2001; Fuchs *et al*, 2004). When departing away from the niche, SCs sense the lack of self-renewal signals or the presence of commitment cues and respond by activating differentiation programs. Likewise, following SC loss, committed cells repopulate the niche and undergo reprogramming into bona fide SCs, a process which depends on an intact niche (van Es *et al*, 2012; Rompolas *et al*, 2013; Nasser *et al*, 2018; Lin *et al*, 2018). To date, however, very little is known about SC – niche crosstalk *in vivo*. A better understanding of SC regulation by the niche is important to clarify mechanisms of SC function under homeostasis, involvement in aging and pathology. Current SC cultures are very limited by the gap in knowledge of niche essential factors. Consequently, primary cultures typically sustain for very short periods, as SCs seemingly lose self-renewal and long-term proliferation capacity when grown outside the niche in the culture dish. This deficit severely halts SC application in regenerative medicine.

SC responses to mechanical cues from various niche components are largely unknown and in recent years starting to be addressed *in vivo* (Bhattacharya *et al*, 2022). In general, few adult SC niches seem to be softer than the differentiation compartments (Eberwein *et al*, 2014; Bornschlögl *et al*, 2016; Monge *et al*, 2017; Gouveia *et al*, 2019). It is hypothesized that SCs can sense the biomechanical properties of their microenvironment and translate this signal into a decision to self-renew or differentiate. A key regulatory step in responses to mechanotransduction pathways involves changes in the localization of the co-transcriptional regulator, YAP, a process that seems to depend on tissue and cell type. In many cases, in response to stiff matrix sensing, the co-transcriptional regulator, YAP, translocates from the cytoplasm to the nucleus and activates mechano-responsive pathways (Dupont *et al*, 2011; Halder *et al*, 2012). However, in a few recent studies, mouse mammary gland stromal fibroblasts (Lerche *et al*, 2020), murine incisor SCs (Lerche *et al*, 2020), and muscle SCs (Eliazer *et al*, 2019) show opposite responses of YAP localization to matrix stiffness or mechanotransduction stimulation by the Rho-A pathway. Another important regulatory step involves Hippo kinases (e.g., LATS1/2 kinases) which can phosphorylate YAP on serine residue 127 (S127), rendering it inactive or prone to degradation (Zhao *et al*, 2010).

The corneal epithelium is excellent *in vivo* model to study the SC niche. The SCs reside in a well-demarcated domain of the limbus, at the ring-shaped zone of the tissue boundary with neighboring conjunctiva, whereas differentiated cells reside in the central cornea that is macroscopically distinguishable. Quiescent and active limbal SC populations function to constantly renew the corneal epithelium under homeostasis and efficiently repair central corneal injury (Cotsarelis *et al*, 1989; Amitai-Lange *et al*, 2015; Di Girolamo *et al*, 2015; Dorà *et al*, 2015; Altshuler *et al*, 2021). Interestingly, limbal SCs (LSCs) are supported by the underlying mesenchymal stromal niche that is less rigid compared to the differentiated zone (Eberwein *et al*, 2014; Gouveia *et al*, 2019). In the corneal epithelial lineage, the expression pattern and function of YAP are unclear. In one study, YAP was found in the nucleus and cytoplasm of murine limbus and cornea (Kasetti *et al*, 2016). Another report linked nuclear YAP with corneal differentiation (Gouveia *et al*, 2019) whereas nuclear YAP was not detected in limbus and cornea in another study (Raghunathan *et al*, 2014).

Here we report that LSCs predominantly express nuclear YAP, which is essential for their functionality. While the limbal SC niche stiffness may be considered relatively stiff for many cell types, the corneal differentiation compartment is much stiffer, and the compartmentalization process involves a distinction between the stiffnesses of these two regions. We propose that matrix rigidity influences YAP activity to regulate the proliferation and wound healing response of LSCs, and the plasticity of committed cells. The higher corneal rigidity increases actomyosin contractility and renders YAP in the cytoplasm, a mechanism that is mediated by activation of SMAD2/3. Inhibiting the process of rigidity sensation or TGF-β pathway significantly improves LSC maintenance and might aid the expansion of LSCs ex-vivo for therapeutic needs.

## Results

### YAP is essential for the maintenance of undifferentiated human LSC state

To characterize YAP’s expression pattern and function, we studied its expression *in vivo* in human tissue sections. Immunohistochemistry indicated that nuclear YAP (nYAP) was predominantly detected in the K15+ SC compartment of the limbus, but not in the corneal center (Fig.1A). To study the role of YAP in LSCs, we established primary human limbal cultures grown in co-culture with mitotically inactive fibroblast feeder cells (Bhattacharya *et al*, 2019). In agreement with *in vivo* data, nYAP was observed in cells expressing the stem/progenitor cell markers K15 and P63 in the colony periphery (Fig. 1B-C, yellow arrows). In line, cytosolic YAP (cYAP) was observed in K15/P63-negative differentiated cells in the center of colonies (Fig. 1B, white arrow). Upon phosphorylation of site S127, YAP becomes cytoplasmic, unable to carry out its putative nuclear co-transcriptional activities (Zheng & Pan, 2019; Ma *et al*, 2019). Specific antibody against the phosphorylated from of YAP (pYAP) confirmed that pYAP is preferentially found in the cytosol of differentiated cells (Fig. 1C, white arrows). This data suggests that nYAP localization and activity is linked with the undifferentiated cell state and that cYAP is linked with cell differentiation. To further confirm this data, we explored YAP expression before and after calcium-induced LSC differentiation (Bhattacharya *et al*, 2019). LSCs were grown for seven days in a defined medium containing 150 μM (low) calcium to minimize differentiation, whereas 1.2 mM (high) calcium was used to induce differentiation. Quantitative real-time polymerase chain reaction (qPCR) and Western blot analyses confirmed successful cell differentiation hallmarked by the reduction in the levels of stem/progenitor cell markers (K15, TP63) and increased expression of differentiation markers (K3 and K12) (Fig. 1D-E). Here too, nYAP was predominantly detected in undifferentiated LSCs, while calcium-driven differentiation enhanced a switch to cYAP (Fig. 1F-G).

**Figure 1.**
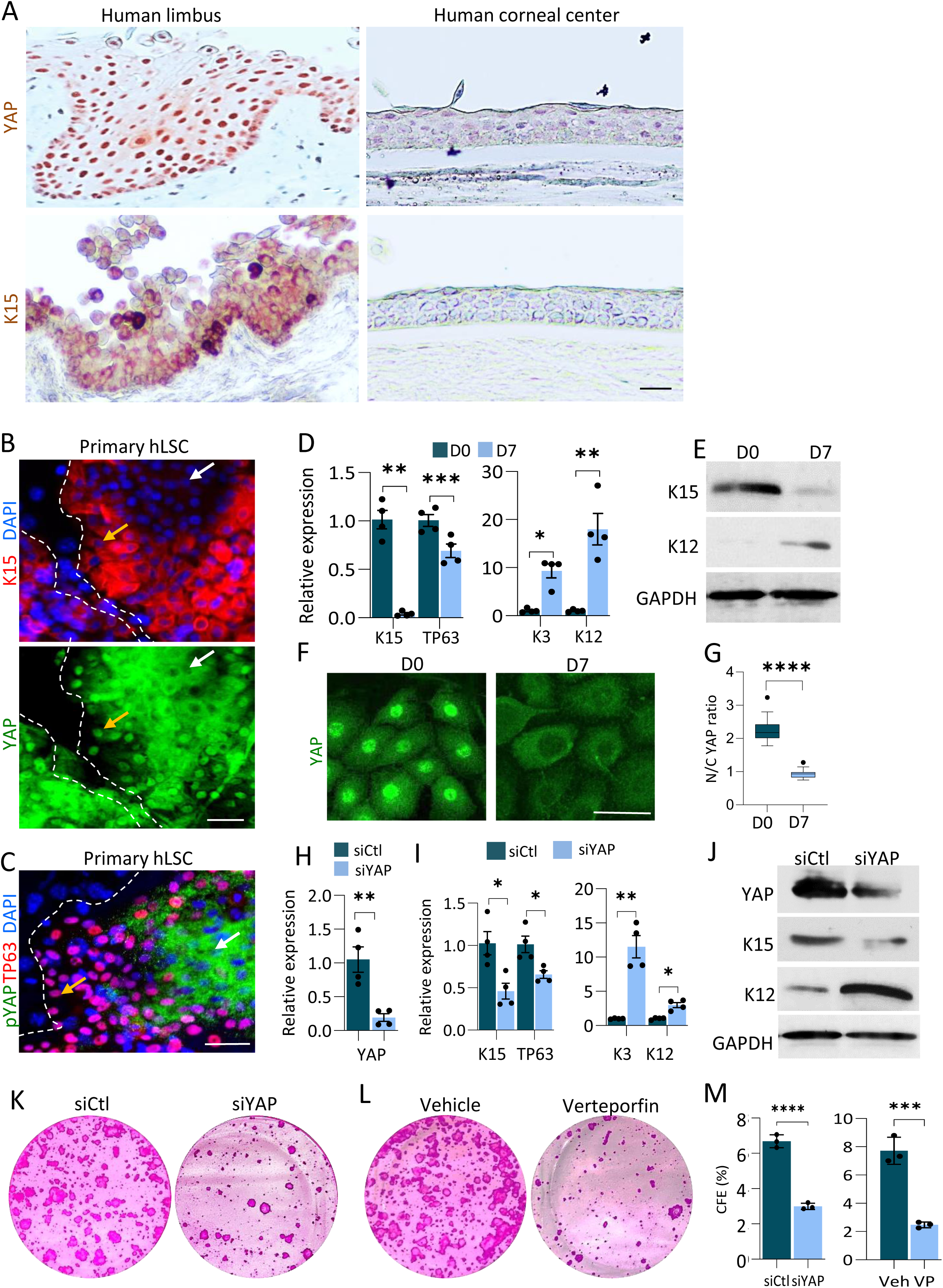
YAP is required to maintain LSC at undifferentiated state *in vitro*. (A) Immunohistochemistry of YAP and K15 was performed on paraffin sections human limbus and cornea. (B-C) Primary human LSCs were co-cultured with NIH-3T3-J2 feeder cells, grown on a plastic plate for 4 days and the expression of the indicated markers was examined by immunostaining. The golden and white arrows indicate SC or differentiated cell regions, respectively. (D-G) Primary human LSCs were grown at low calcium (Day 0) or induced to differentiate in high calcium (Day 7) and the expression of the indicated markers was tested by quantitative real time PCR (qPCR) (D), Western blot (E) or immunostaining (F) that was further quantified for mean nuclear-to-cytoplasmic YAP intensity ratio (G). (H-J) Primary LSCs were transfected with esiRNA against YAP (siYAP) or control esiRNA (siCtl) and 48-72 hours later, the expression of YAP and the indicated markers was examined by qPCR (H, I) and Western blot (J). (K-M) Primary LSCs were transfected with siYAP or siCtl or treated with Verteporfin or vehicle, and 48-72 hours later, cells were subjected to a clonogenicity test (K, L) and the number of the colony was quantified (M). The qPCR data were normalized to the housekeeping gene (mean ± standard error of mean, n=4 biological replicates) as a fold increase compared to the control sample and mean nuclear-to-cytoplasmic YAP intensity ratio (G) is shown by the Tukey box-and-whisker plot. Data represents ≥3 biological replicates. Statistical significance was assessed by t-test (*, *p* < .05; **, *p* < .01; ***, *p* < .001, ****, *p* < .0001). Nuclei were detected by DAPI counterstaining. Scale bars are 50µm.

To investigate the influence of YAP on stemness, we performed knockdown experiments using endonuclease-digested silencing RNA (known as esiRNA) that are considered to have fewer off targets and higher efficiency (Jeric *et al*, 2016; Price *et al*, 2021). YAP repression by esiRNA was efficient and resulted in a significant reduction in expression of the stem/progenitor cell markers (K15, TP63) and an increase in the expression of the differentiation markers (K3, K12) (Fig. 1H-J). To further exclude the possibility of non-specific reagent activity, knockdown was performed with 2 conventional previously validated silencing RNAs against YAP (siYAP) (Dupont *et al*, 2011) or with the pharmacological inhibitor of YAP, Verteporfin (Ning *et al*, 2021; McGinn *et al*, 2021; Jiang *et al*, 2021). In agreement, both siRNA repression (S1A-B) and pharmacology inhibition (Fig. S1C) efficiently repressed YAP and the stemness phenotype. We next explored the impact of YAP knockdown by esiRNA or inhibition by Verteporfin treatment on the ability of LSCs to form colonies when grown at low density in vitro, a key hallmark of LSCs. To that end, cells were seeded at clonal density and allowed to expand for 2 weeks in co-culture with feeder cells. As shown in Fig. 1K-M, colony-forming efficiency was drastically affected by YAP knockdown or Verteporfin treatment, suggesting that YAP plays a crucial role in preserving long-term LSC proliferation.

### YAP inhibition perturbs LSC function *in vivo*

To validate the data and delineate YAP’s underlying role in LSC regulation *in vivo*, we characterized the expression pattern of YAP in murine. Gene expression patterns and lineage tracing experiments implies that restriction of SCs to the limbus and renewal of the cornea by LSCs is established around post-natal day 15 (P15) and that the cornea fully matures by P60 (Collinson *et al*, 2002; Dorà *et al*, 2015; Richardson *et al*, 2017). Interestingly, in the immature P15 corneas, the LSC reporter transgene K15-GFP (Fig. 2A) and K15 protein (Fig. 2B) were detected in a punctuated pattern throughout both the limbal and central corneal epithelium, suggesting that at this stage SC are located in both limbus and cornea zones. By P60, however, the labeling pattern of K15-GFP and K15 (Fig. 2A-B) became sharply constrained to the limbus. Co-immunofluorescent staining confirmed that nYAP is expressed by K15+ LSCs (Fig. S2A). To furthermore, reaffirm the conclusion that YAP is correlated with stemness, To furthermore validate the link between active nYAP and LSCs, we investigated the expression pattern of the Hippo pathway kinases, LATS1/2, which can phosphorylate and inactivate YAP. Staining with antibodies that specifically recognize the active phosphorylated LATS1/2 (pLATS1/2) and pYAP, confirmed that pLATS1/2 and pYAP are predominantly found in the corneal differentiation compartment at P60 (Fig. S2B-C). This observation suggests that in the cornea, LATS1/2 kinases become activated in the differentiated corneal cells and consequently phosphorylate YAP, thereby preventing its nuclear translocation.

**Figure 2.**
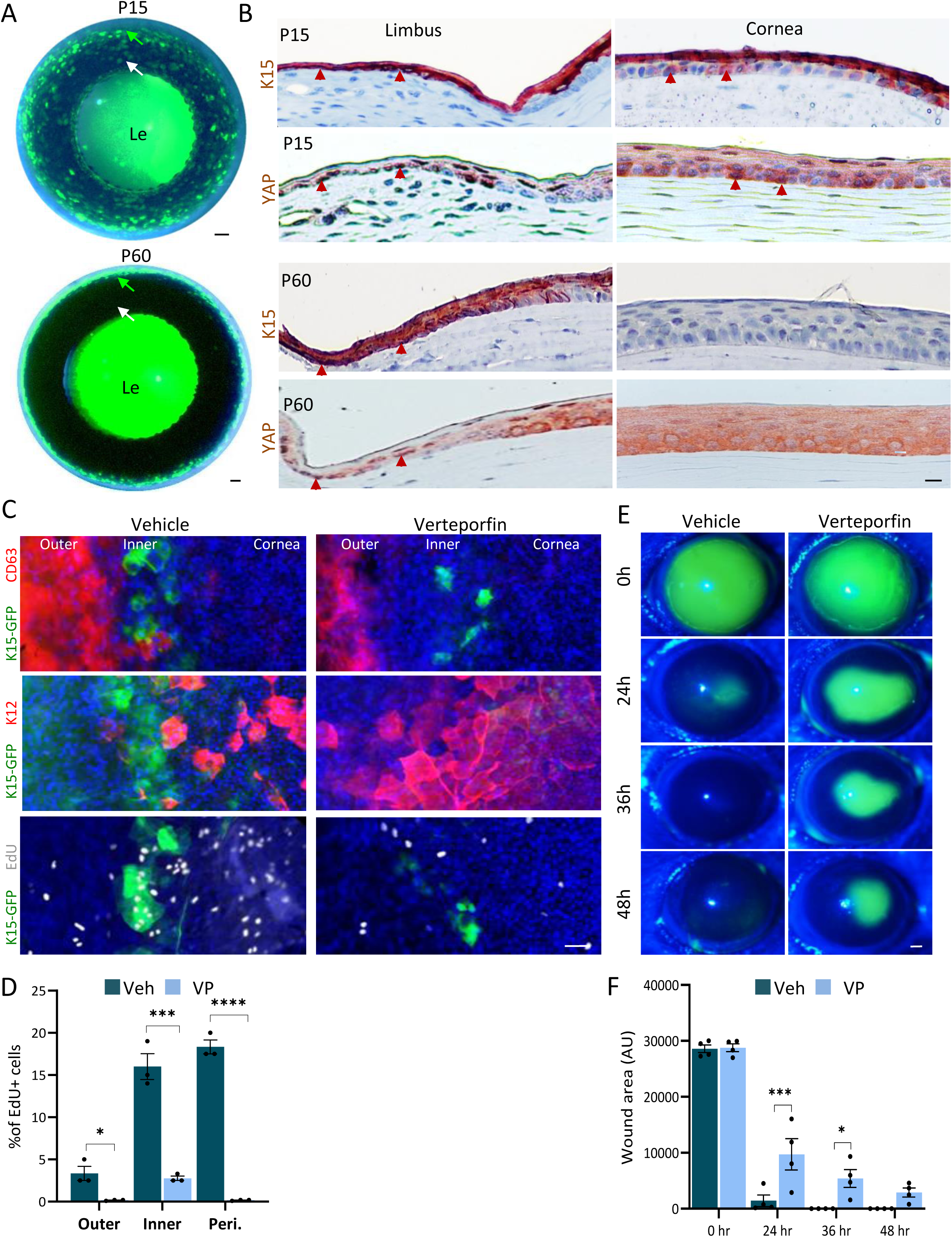
YAP inhibitor perturbs LSC function *in vivo*. (A) K15-GFP transgenic animals were sacrificed at P15 or P60 and the whole eye merged bright field and fluorescent images are shown. Green and white arrows indicate the limbus and cornea region, respectively. (B) Immunohistochemistry was performed on paraffin sections of the P15 or P60 wild-type mouse cornea using the indicated antibodies. Arrowheads indicate nuclear YAP or K15 expression (C, D) Daily sub-conjunctival injections (20µl) of Verteporfin (20µM) or vehicle (control) were performed for 4-days to adult K15-GFP transgenic mice. EdU was injected 6-hours before eyes were enucleated and prepared for wholemount immunostaining. CD63+ outer quiescent LSCs, K15-GFP+ inner active LSCs or K12+ corneal epithelial differentiated cells are shown, among them, EdU+ cells that are in mitosis and were quantified in (D) (n=3 biological replicates). (E-F) The corneal epithelium of P60 mice was surgically removed using Algerbrush and animal were treated with Verteporfin (20 µM) or vehicle. At indicated time points, fluorescein dye staining was pictured (E) to follow epithelial healing and quantification is shown in (F) (n=4 biological replicates). All data represents ≥3 biological replicates and are represented as mean ± standard error of mean. Nuclei were detected by DAPI counterstaining. Scale bars are 500 µm (A) and the rest are 50µm. Abbreviation: Le, Lens.

To test YAP involvement with LSC function *in vivo*, we performed sub-conjunctival injections (15 µl) of Verteporfin (20 µM) or vehicle (as control) once a day for four days. Analysis of cell and tissue morphology and apoptosis confirmed that the treatments with Verteporfin did not trigger cell loss or apoptosis (Fig. S3). Recent studies indicated that the limbus contains 2 clear LSC domains, the “outer” LSC zone hosts LSC population that serve as a reservoir for wound healing whereas the “inner” LSCs actively renew the cornea (Altshuler *et al*, 2021; Farrelly *et al*, 2021). As shown in Fig. 2C, the YAP inhibitor, Verteporfin (VP), repressed markers of the outer (CD63) and inner (K15-GFP) LSCs and increased the differentiation marker, K12, in the limbus. Moreover, Verteporfin treatment decreased cell proliferation, as evidenced by lower incorporation of the nucleotide analogue 5-Ethynyl-2’-deoxyuridine (EdU) into the cells’ DNA (Fig. 2C-D). To further explore the link between YAP and LSC functionality, we performed a large (2 mm) corneal epithelial debridement and followed epithelial healing with a fluorescein dye penetration test. As shown in Fig. 2E-F, Verteporfin significantly delayed wound healing, suggesting that YAP regulates LSC activity *in vivo*. Altogether, these data suggest that YAP regulates SC phenotype and cornea integrity *in vivo*.

### YAP inhibitor perturbs niche mediated dedifferentiation of corneal committed cells

Catastrophic loss of the entire LSC population can be recovered owing to the high plasticity of corneal epithelial committed cells (Nasser *et al*, 2018). The latter cells repopulate the denuded limbus, and in case the niche is intact, they dedifferentiate into K15-GFP+ LSC-like cells and the corneal tissue integrity is maintained for many months. However, the mechanism regulating this process by the niche is not clear. Next, we explored if along with LSC maintenance, YAP is also required for reprogramming of corneal committed cells. To explore this hypothesis, the entire limbal epithelium (including marginal conjunctival and bordering peripheral corneal epithelium) was surgically removed, rendering the cornea deprived of LSCs (as shown by Nasser *et al*, 2018). Complete repopulation of the limbal epithelium by corneal committed cells expressing nYAP was already evident by day one post-injury, and the signal became even more pronounced and widespread over time (Fig. S4), suggesting that YAP might be involved in the dedifferentiation process. To check this possibility and properly follow regeneration and the dedifferentiation processes in real-time, we used the triple transgenic K14-Cre^ERT2^; R26R-Brainbow^2.1^; K15-GFP mice (Nasser *et al*, 2018). In this multi-color “Confetti” lineage tracing system, transient exposure of 2/3-months old transgenic animals to tamoxifen (3-4 days injections) induces the stochastic and irreversible labeling of K14+ limbal/corneal basal cells with one out of four fluorescent protein-coding genes (i.e. cytoplasmic red (RFP) or yellow (YFP), membrane cyan (CFP) or nuclear green (GFP)) (Amitai-Lange *et al*, 2015; Nasser *et al*, 2018; Altshuler *et al*, 2021) (Fig. 3A-C). Under homeostasis, the centripetal renewal of the corneal epithelium by Confetti+ LSCs can be visualized by intravital microscopy over time. Large Confetti+ limbal radial clones that emerged from the K15-GFP+ inner limbus were evident four months post-induction (Fig. 3D, unwounded, yellow arrows mark double positive cells). As illustrated in Fig. 3C, surgical limbal epithelial removal (LER) was performed (see LER in Fig. 3D) and tissue regeneration was traced following subconjunctival injection of Verteporfin or vehicle (as control). Injections were performed every other day until 10 days when recovery of K15-GFP signal peaked (see D10 in Fig. 3D), and at this point, treatment was ceased to avoid the potentially harmful effects of multiple eye injections. In all treatments, by day 1 (D1) post LER, stripes of corneal committed cells successfully repopulated the debrided limbal epithelium (Fig. 3D). Intriguingly, however, the K15-GFP recovery was markedly affected by Verteporfin treatment (Fig. 3D, see D7-15). Moreover, on D15 (5-days post last injection), the expression of LSC markers (K15-GFP, CD63 and GPHA2) was poorly revived in the limbus following Verteporfin treatment (Fig. 3D-E). This data suggests that YAP plays a vital role in the reprogramming of corneal committed cells into LSC-like cells.

**Figure 3.**
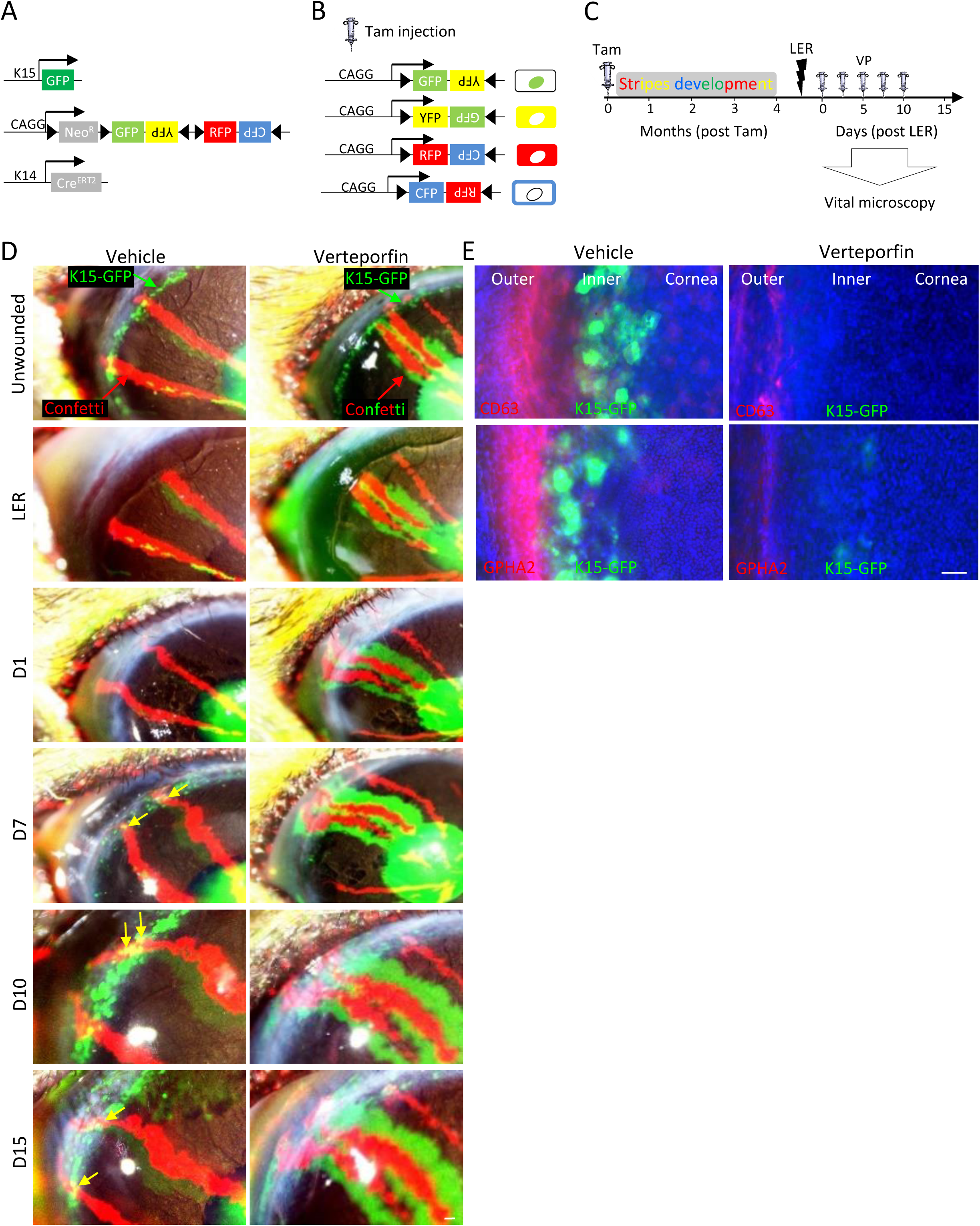
YAP inhibitor perturbs niche mediated dedifferentiation. (A) Schematic illustration of triple transgenic animals (K15-GFP; Brainbow^2.1^; K14-Cre^ERT2^) used, (B) examples of potential the reorganization of Brainbow cassette after tamoxifen induction. (C-E) As illustrated in (C), 2-3 moth old triple transgenic animals were injected with tamoxifen (Tam) to induce the random and irreversible expression of Confetti reporters (B). (C-E) Four-months post Tam induction, fully developed Confetti+ (RFP+) limbal radial stripes were evident (see unwounded), then, the limbal epithelium was surgically removed (see LER) and mice were treated Verteporfin (VP, 20 µM) or control (Vehicle). For accurate clonal tracking over time, eyes with 1–2 RFP^+^ stripes were pictured (n = 5 biological replicates) in live animals (D). In vehicle treated mice, Confetti + (RFP^+^) corneal-committed cells repaired the denuded limbus by day 1 (D1) post LER and re-expressed K15-GFP by D10. In Verteporfin treated cornea, the recovery of K15-GFP was negligible. (E) On D15 post LER, eyes were enucleated and wholemount immunostaining for markers of the quiescent outer (GPHA2, CD63) LSCs or inner active (K15-GFP) LSCs is shown. Data represent 5 biological replicates. Nuclei were detected by DAPI counterstaining. Scale bars are 50µm.

### Manipulation of stiffness impaired SC function and YAP localization *in vivo*

Very little is known about the limbal niche formation and involvement of biomechanical cues in this process. Previous measurements of the rigidity of the human cornea indicated that the limbus is a relatively rigid tissue (8 kPa), however, the corneal center is much stiffer (20 kPa) (Eberwein *et al*, 2014). But the rigidity in the murine limbus, which differs in anatomy from that of the human limbus, has never been tested. To analyze the underlying rigidity that murine basal limbal/corneal layer cells may sense, the entire limbal/corneal epithelial layer was removed by ethylenediaminetetraacetic acid (EDTA) treatment, and the rigidity of the denuded surface was tested by atomic force microscopy (AFM). At P15, the stiffness of the murine cornea and limbus were comparable (∼1 kPa) where SCs were uniformly scattered through the cornea (Fig. 4A). In the mature P60 murine, the limbus, although not as stiff as the human limbus, is a relatively rigid tissue (∼2 kPa) while the corneal differentiation compartment is significantly stiffer (∼7 kPa) (Fig. 4A). Together with the confinement of SCs to the limbus and the nYAP phenotype observed in that region (Fig. 2A-B), this data suggests that matrix stiffening might play an important role in LSC regulation.

**Figure 4.**
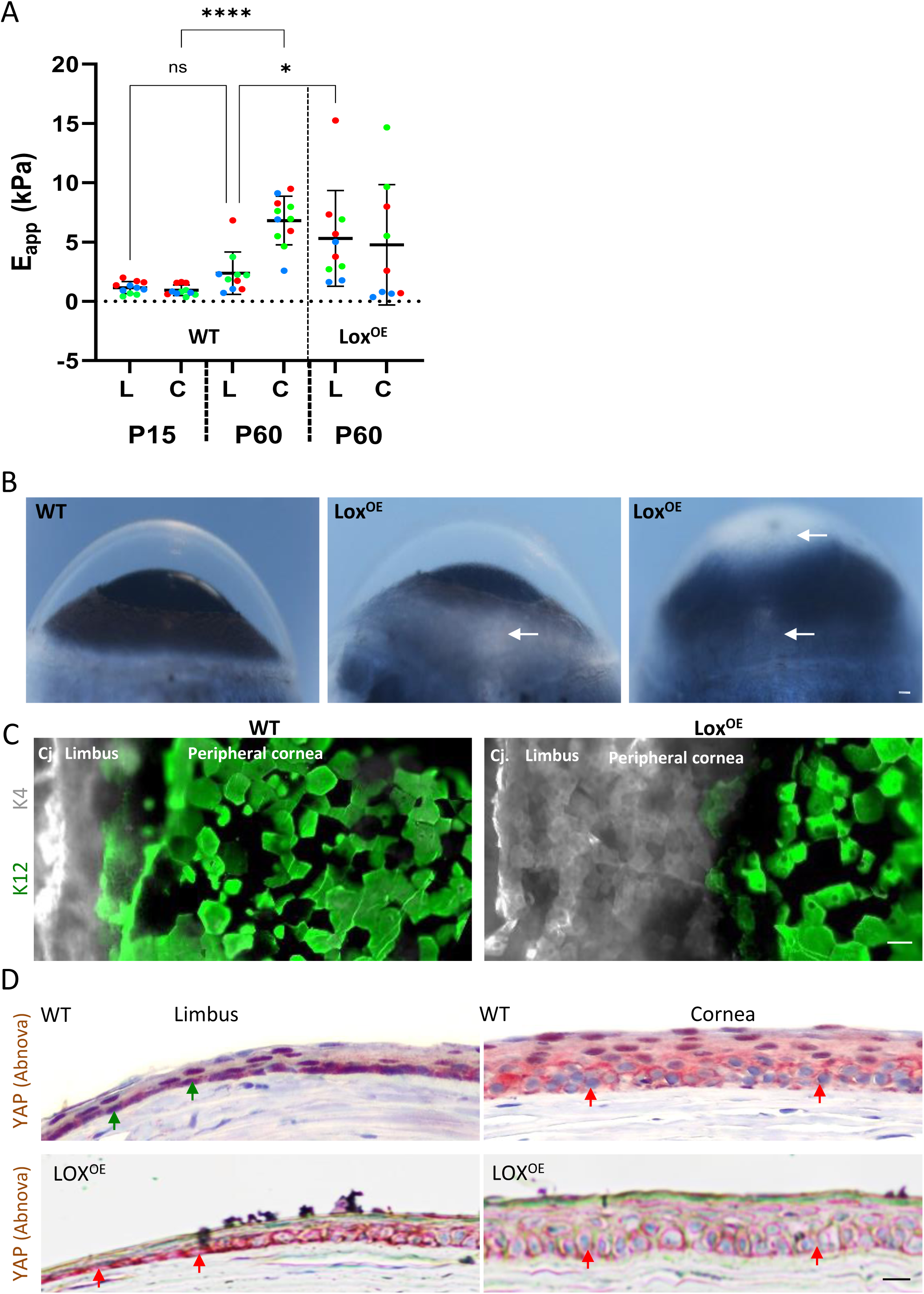
Manipulation of niche rigidity impairs SC phenotype and corneal integrity. (A) Atomic force microscopy measurement of the rigidity of the limbus (L) and the cornea (C) in wild type (WT) and Lox transgenic (Lox^OE^) mice at post-natal day 15 mice (P15) and P60 (the higher the Eapp the stiffer the tissue). Mean +/- SD of the apparent Elastic modulus (in kPa). The plot shows the merge of three experiments performed on as many independent samples, using the same atomic force microscopy setup (deflection sensibility and cantilever spring constant). (B) Bright field binocular images of the eyes of the indicated genotypes are shown. Lox^OE^ corneas often display peripheral and/or central opacification (arrows). (C) Wholemount immunostaining for conjunctiva (K4) and cornea (K12) markers of the indicated genotypes. (D) Immunohistochemistry using anti-YAP (Abnova) antibody on paraffin sections of the P60 cornea of the indicated genotype. Green arrows indicate nYAP and red arrows indicate cYAP. Data represents ≥3 biological replicates. Scale bars are 50µm. Abbreviation: Cj. Conjunctiva.

To further explore whether matrix stiffening alone can induce differentiation, we investigated the corneas of adult transgenic animals that over-expressed the enzyme Lysyl oxidase (Lox) (Gabay Yehezkely *et al*, 2020). Lox catalyzes covalent crosslinking of collagen and elastin in the extracellular matrix (ECM), thereby increasing tissue stiffness (Levental *et al*, 2009; Baker *et al*, 2013; Vallet & Ricard-Blum, 2019). AFM measurements revealed a ∼2-fold increase in the stiffness of adult limbus of transgenic Lox (Lox^OE^) animals (Fig. 4A). Moreover, Lox^OE^ corneas often displayed typical hallmarks of a clinical entity known as “LSC deficiency” (Huang & Tseng, 1991), including corneal opacification (Fig. 4B) coupled with the abnormal presence of cells that express conjunctival cell markers in the cornea (Fig. 4C), suggesting for conjunctival cell invasion into the cornea. In line with enhanced niche stiffness, Lox^OE^ mice expressed cYAP in both the limbus and cornea (Fig. 4D). This data implies that LSCs can sense the stiffness of the niche matrix and that this biomechanical signal affects YAP localization and LSC function *in vivo*.

### Rigidity-induced adhesion and actomyosin contractility enhances cell differentiation through YAP inactivation

The association between the relatively low limbal rigidity, nYAP and stemness (Fig. 1, 4D), suggests that corneal rigidity reduces nYAP to enhance differentiation. These relationships are inconsistent with many studies that linked nYAP with stiffer matrices and/or with activation of the RhoA pathway (Dupont *et al*, 2011; Elosegui-Artola *et al*, 2016; Totaro *et al*, 2017). However, other studies have recently reported that lower rigidity or inhibition of the Rho-A pathway enhanced nYAP in mammary gland stromal fibroblasts (Lerche *et al*, 2020), mouse incisor SCs (Otsu *et al*, 2021) and muscle SCs (Eliazer *et al*, 2019), suggesting that cell context-specific factors modulate the response of YAP to substrate rigidity.

As a control experiment, we first used two fibroblast cell lines, mouse embryonic fibroblasts (MEFs) and WI-38, which were expected to display “standard” YAP responses to rigidity. We grew both cell lines on silicone gels of substrate stiffness ranging from 0.25 to 35 kPa and found a positive correlation between rigidity and nYAP (Fig. S5A-B). To further test the effect of substrate rigidity on YAP localization, primary human LSCs were cultivated for 4-days on silicone gels that recapitulate the rigidity of the human limbus (8 kPa, referred to hereafter as “limbal rigidity”) or cornea (20 kPa, “corneal rigidity”) (Eberwein *et al*, 2014). In line with a previous report (Gouveia *et al*, 2019), growth on corneal rigidity-like gels decreased the expression of stem/progenitor cell markers (K15, TP63) and increased differentiation markers (K3, K12) (Fig. 5A-B, S5C-D), as compared to control gels (limbal rigidity). In agreement with the *in vivo* data in mice (Fig. 2B, Fig. S2B-C), human LSCs grown on the corneal rigidity for 4-days displayed much higher levels of pLATS1/2 and cYAP (Fig. 5C-D). The stiffer substrate enhanced focal adhesion maturation as evidenced by Vinculin staining (Fig. 5C, S6A) and cell spreading (Fig. S6C), altogether suggesting that LSC differentiation is linked to activation of the mechanotransduction pathways. Next, we tested whether the rigidity-dependent YAP translocation and cell differentiation require actomyosin contractility. To this end, cells were seeded on matrices with corneal stiffness and actomyosin-mediated forces were inhibited by Blebbistatin (20 μM) for 4-days. Indeed, Blebbistatin attenuated the stiffness-induced transition to mainly cYAP (Fig. 5E-F), attenuated cell differentiation (Fig. 5E, G) and cell spreading (Fig. S6B). Collectively, these data suggest that stiffness-induced differentiation is controlled by LATS1/2 mediated cYAP localization through actomyosin contractility.

**Figure 5.**
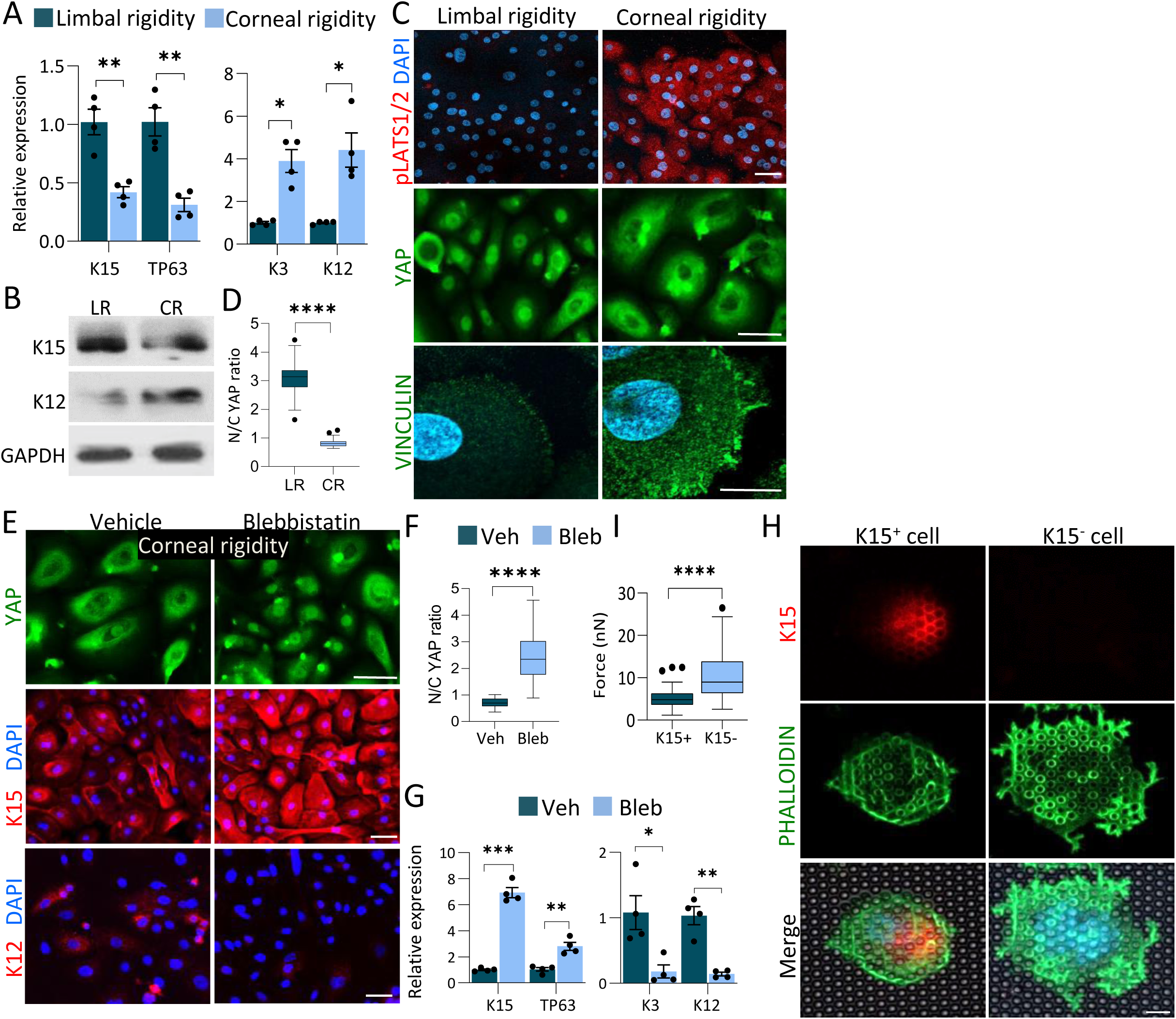
Corneal rigidity induces differentiation and inhibits nuclear localization of YAP. Primary human LSCs were grown on a silicone substrate that mimics human limbus (LR, 8kPa) or corneal (CR, 20kPa) rigidity that is coated with fibronectin and cultured for 4 days. Expression of SC (K15, TP63) or differentiation (K3, K12) markers was tested by quantitative real time PCR (qPCR) (A) or Western blot (B) or immunostaining with the indicated antibodies (C) and mean nuclear-to-cytoplasmic YAP intensity ratio was quantified (D). (E-G) Primary LSCs were grown on silicone substrate that mimics corneal rigidity, were cultured for 4 days with Blebbistatin (Bleb) or Vehicle (Veh) and the expression of the indicated markers was tested by immunostaining (E) and mean nuclear-to-cytoplasmic YAP intensity ratio was quantified (F) or the expression of the indicated genes was tested by qPCR (G). (H-I) Primary human LSCs were seeded on pillars that mimic limbal or corneal rigidity (right) overnight. Immunostaining of K15 and F-actin (phalloidin) is shown in (H) and measurements of forces generated by K15-positive and K15- negative cells (undifferentiated and differentiated; see Methods) shown in (I); n=44 and 52 cells respectively from three independent experiments. Mean nuclear-to-cytoplasmic YAP intensity ratio in (D, F) is shown by Tukey box-and-whisker. Force generated by LSCs is shown in (H) by Tukey box-and- whisker plot followed by Mann Whitney test. The qPCR data were normalized to the housekeeping gene and is presented (mean ± standard error of mean, n=4 biological replicates) as fold increase compared to control sample and statistical analysis was performed by t-test (*, p < .05; **, p < .01; ***, p < .001, ****, p < .0001). Data represents ≥3 biological replicates. Nuclei were detected by DAPI counterstaining. Scale bars are 5 µm (H) and the rest are 50µm.

We next turned to characterize the contractile forces that the LSCs produce for mechanosensing. To that end, we used arrays of fibronectin-coated elastic polydimethylsiloxane (PDMS) pillars with limbal or corneal rigidity (Wolfenson *et al*, 2016; Meacci *et al*, 2016). After overnight incubation, the cell size was significantly smaller on the pillars with limbal rigidity and a clear inverse correlation between cell area and K15 expression was evident (Fig. S6C-D). Moreover, larger K15-negative differentiated cells typically displayed filamentous actin (F-actin) distributed as rings around individual pillars at the cell edge; in contrast, smaller K15+ cells typically showed much fewer F-actin rings, and rarely at the edge (Fig. 5H). F-actin around pillars is positively correlated with the level of the force on the pillars (Feld *et al*, 2020); indeed, the cells that lacked K15 expression generated on average almost 2-fold higher forces on individual pillars compared to cells that had high K15 expression (Fig. 5I). This observation is in line with the formation of larger adhesions on the stiff matrix (Fig. 5C, S6A), which support the transmission of higher contractile forces. Altogether, these data suggest that matrices with limbal rigidity support the formation of small adhesions and weaker contractile forces that render low LATS1/2 activity and consequently favor stemness through maintaining nYAP.

### SMAD2/3 represses nYAP and mediates stiffness-induced differentiation

The human limbus is relatively stiff (8 kPa) whereas the cornea is very stiff (20 kPa). We, therefore, suspected that the very stiff corneal rigidity stimulates another signaling pathway that overrides the “standard” mechanotransduced nYAP location, thereby leading to cYAP localization. Recent studies have shown that SMAD2/3 inhibition attenuates epithelial cell differentiation and that there is synergy between myosin II and TGF-β in the regulation of SC differentiation (Mou *et al*, 2016; Zhang *et al*, 2018). Typically, active phosphorylated SMAD2/3 proteins translocate to the nucleus, where they regulate transcription (Massagué, 1998). Indeed, nuclear SMAD2/3 were preferentially detected following calcium-induced differentiation of LSCs on plastic (Fig. S7) or following either 4-days growth on corneal rigidity. Treatment with Rho Activator II (Rho-Act; 1 µg/ml) which is known to enhance the mechanotransduction pathway (Lessey *et al*, 2012; Dias Gomes *et al*, 2019; Burridge *et al*, 2019) was sufficient to induce nuclear SMAD2/3 (Fig. 6A-B).

**Figure 6.**
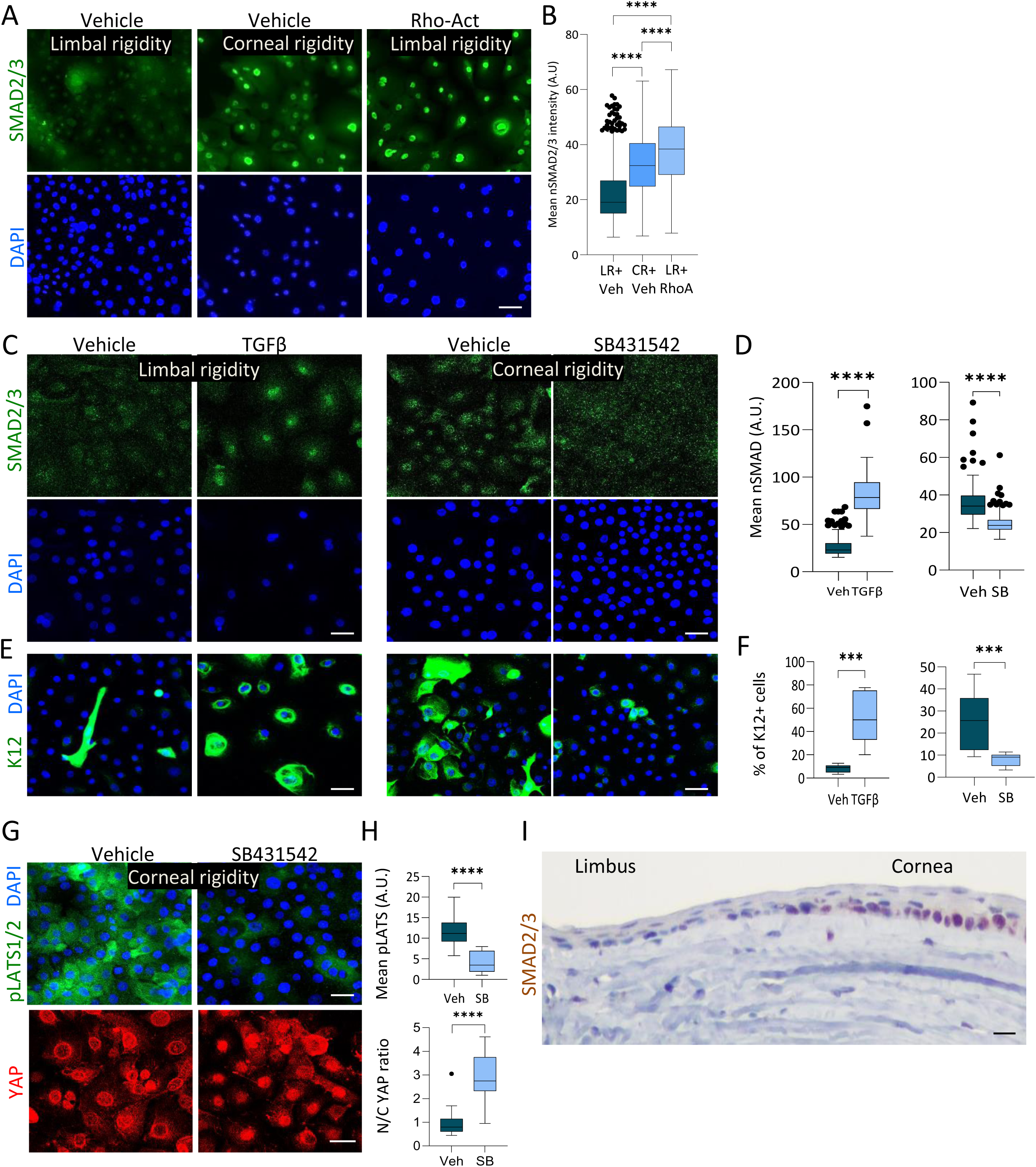
Stiffness-induced differentiation is mediated by SMAD2/3. (A-H) Primary human LSCs were treated with Rho-Activator (Rho-Act) or Verteporfin (VP) for 4 days whereas treatment with TGFβ ligand or TGFβ pathway inhibitor (SB431542), or with control vehicle for 2 days. Cells were then fixed and immunostained with the indicated antibodies (A, C, E, G) and quantification of mean nuclear SMAD2/3 (nSMAD)(D), % of K12+ cells (E), or mean nuclear-to-cytoplasmic YAP intensity ratio and mean pLATS1/2 intensity is shown (H). (L) Immunohistochemistry of SMAD2/3 was performed on paraffin sections of the P60 mouse cornea. The regions of the limbus and corneal periphery are shown. Data represents ≥3 biological replicates. Nuclei were detected by DAPI counterstaining and scale bars are 50µm. Mean nuclear-to-cytoplasmic YAP intensity ratio, mean pLATS1/2 intensity, and mean nuclear SMAD2/3 intensity is shown by the Tukey box-and-whisker plot followed by Mann Whitney test. % of K12 positive cells is shown by Tukey box-and-whisker plot followed by t-test with Welch’s correction (*, p < .05; **, p < .01; ***, p < .001, ****, p < .0001).

Activation of SMAD2/3 with recombinant TGFβ on limbal rigidity sufficiently induced cell differentiation whereas inhibition of the TGFβ pathway by SB431542 on corneal rigidity reduced nuclear SMAD2/3 and inhibited differentiation (Fig. 6C-F). Importantly, the TGFβ pathway inhibitor reduced pLATS1/2 and enhanced nYAP (Fig. 6G-H), suggesting an inverse regulation between TGFβ/SMAD2/3 and YAP. Finally, immunohistochemistry demonstrated that the nuclear SMAD2/3 signal is absent in the limbus, whereas it is prominent in the murine cornea *in vivo* (Fig. 6I). Altogether, these data suggest that the very stiff corneal rigidity induces SMAD2/3-mediated repression of YAP to induce differentiation.

### Inhibition of mechanosensing and TGFβ pathway attenuates LSC differentiation on plastic

Finally, our results led us to consider the fact that plastic dishes that possess extremely high stiffness (Giga Pascals) are widely used to grow SCs for research and cell therapy. Therefore, we examined the impact of inhibition of mechanosensing by Blebbistatin or inhibition of the TGFβ pathway by SB431542 on plastic dishes. Primary LSCs were cultivated with each inhibitor for 4-days before harvesting and analysis. Treatment with either of the inhibitors resulted in enhanced expression of LSC markers, and reduced differentiation gene expression by immunostaining (Fig. 7A-B) and qPCR (Fig. 7C-D). Moreover, these inhibitors significantly augmented colony-forming capacity (Fig. 7E-H). Taken together, we conclude that SC culture on materials that extensively differ in the biomechanical properties of the native niche may affect SC self-renewal and long-term proliferative potential, and manipulating mechanosensing pathways may, at least partially, rescue this undesired effect.

**Figure 7.**
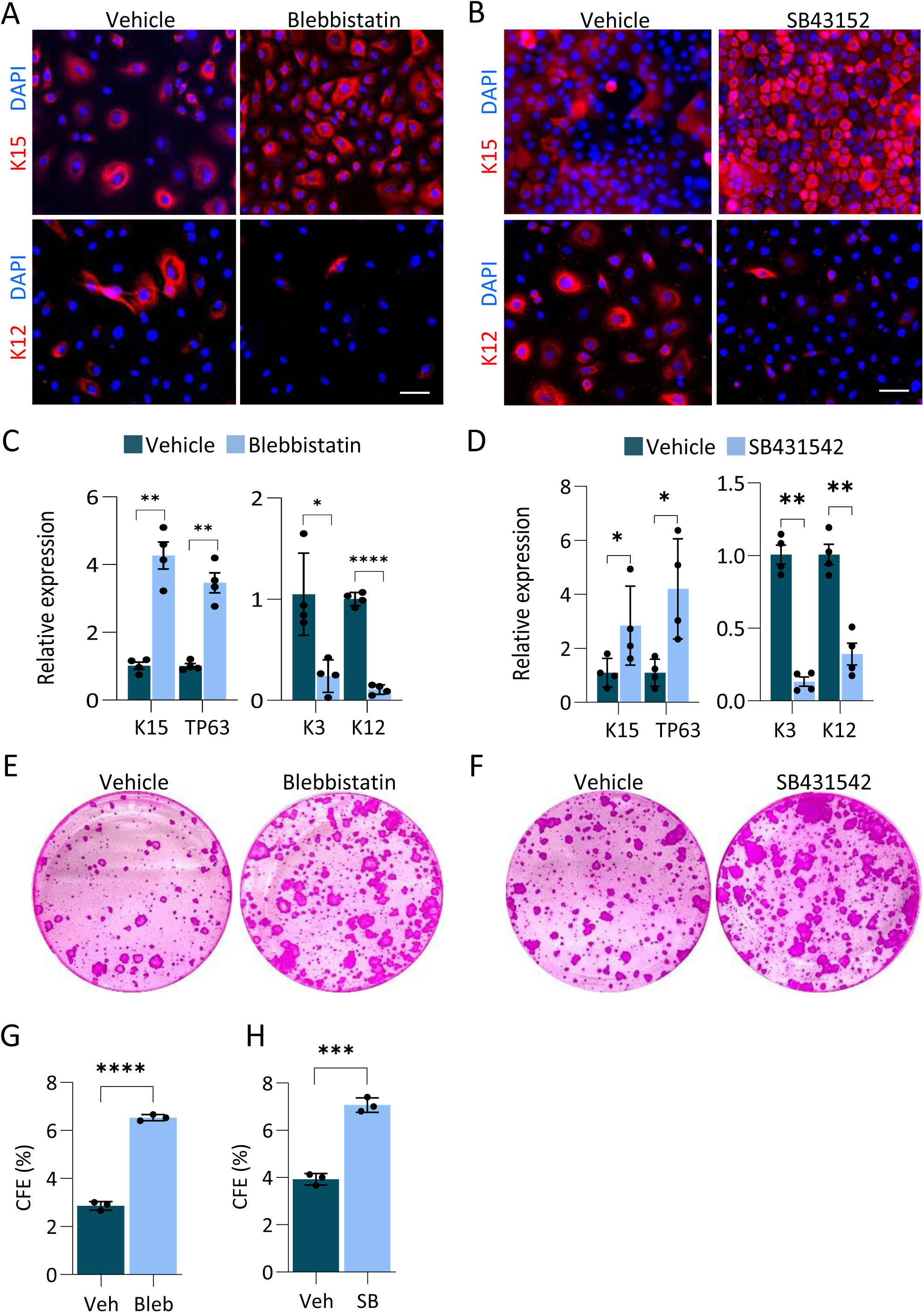
Inhibition of actomyosin contractility or TGFβ pathway attenuates LSC differentiation on plastic. (A-H) Primary human LSCs were grown on plastic and treated with Blebbistatin (Bleb) or TGFβ pathway inhibitor (SB43152) or vehicle (control) for 4 days and the expression of the indicated markers was tested by immunostaining (A-B), quantitative real time PCR analysis (C-D) or cells were subjected to clonogenicity assay and colonies were visualized by Rhodamine stain (E-F) and the number of the colony was quantified (G-H). Real-time data was normalized to the housekeeping gene and is presented (mean ± standard error of mean, n=4 biological replicates) as fold increase compared to control sample and statistical analysis was performed by t-test (*, p < .05; **, p < .01; ***, p < .001, ****, p < .0001). Data represents 3 biological replicates. Nuclei were detected by DAPI counterstaining. Scale bars are 50µm.

## Discussion

SCs can sense the external environment and respond by making cell fate decisions to self-renew or differentiate. A relatively diminutive movement away from the niche to the differentiation compartment (e.g. of 20-30 microns) could sufficiently induce the commitment to the differentiation of Lgr5-GFP+ gut (Barker *et al*, 2007), K15-GFP+ hair follicle and limbal SCs (Morris *et al*, 2004; Nasser *et al*, 2018; Altshuler *et al*, 2021). The importance of the niche in reprogramming is also striking. In response to SC depletion, epithelial cells that have already undergone commitment to differentiation, heal the affected niche, which stimulates their dedifferentiation into SC-like cells (Blanpain & Fuchs, 2014; Tetteh *et al*, 2016; Nasser *et al*, 2018; Lin *et al*, 2018). Mechanotransduction pathways and/or YAP may be involved in this unexpected cell plasticity (Yimlamai *et al*, 2014). To “read” and respond to the rigidity of the underlying matrix, SCs continuously probe the microenvironment and especially the underlying ECM by applying forces, engaging the Rho/actomyosin mechanotransduction pathways.

The influence of YAP on differentiation appears to be cell context-dependent. YAP plays a positive role in the maintenance of adult SCs (Camargo *et al*, 2007; Schlegelmilch *et al*, 2011; Zhang *et al*, 2011; Judson *et al*, 2012; Han *et al*, 2015; Totaro *et al*, 2017; Walko *et al*, 2017) in some cell types, whereas YAP activity was linked with differentiated cells grown on stiff matrices (Dupont *et al*, 2011; Elosegui-Artola *et al*, 2016; Totaro *et al*, 2017); in those cases, nYAP was correlated with adhesion growth (Elosegui-Artola *et al*, 2016) and large cell areas (Dupont *et al*, 2011).

In line with a previous report (Gouveia *et al*, 2019), we observed that the high rigidity of the cornea induces LSC differentiation. The present study, however, supports a role for YAP in the maintenance of an undifferentiated state. The inhibition of nYAP in LSCs is induced by seeding LSCs on a substrate with corneal stiffness, or by TGFβ pathway induction (or by siRNAs or Verteporfin), all of which induced cell differentiation. The role of YAP in controlling stemness is supported by the observation that (i) nYAP was mainly inspected in LSCs, (ii) YAP inhibition by siRNAs or pharmacological inhibitor enhanced differentiation phenotype, (iii) attenuated LSC function and (iv) wound healing response. Theoretically, this could be a secondary effect to the decrease in cell density that accompanies cell differentiation in vitro. However, since comparable effects were found in the *in vivo* settings, where cell densities have not been changed, it is less likely to be the case.

YAP’s importance in the eye and corneal development was evident as single-copy deletion of the *Yap1* gene in mice induced complex ocular abnormalities, including microphthalmia, thinner Descemet’s membrane, and corneal fibrosis, the latter phenotype being reminiscent of LSC deficiency (Kim *et al*, 2019). The coincided stiffening and differentiation of the cornea compartment, imply that mechanosensing processes play a role in tissue development. To further dissect the role of YAP in corneal development, homeostasis and regeneration, future studies should conditionally ablate *Yap1* and YAP-related genes, in a tissue and stage-specific manner.

According to the current dogma, YAP is typically bound to TAZ and both YAP/TAZ cannot bind to the DNA by themselves, they require additional factors to exert their function on gene expression. Several transcription factors have been indicated to be YAP targets, including P73, the ErbB4 cytoplasmic domain, and TEAD (Vassilev *et al*, 2001; Komuro *et al*, 2003; Basu *et al*, 2003). Particularly of interest, YAP has been shown to directly bind and regulate P63 expression in skin keratinocyte SCs (Tomlinson *et al*, 2010) and adult lung basal SCs (Zhao *et al*, 2014), and is implicated in regulating the transcription of SOX2 (Bora-Singhal *et al*, 2015). Since SOX2 and P63 interact and control LSC function (Bhattacharya *et al*, 2019), YAP may bind to P63/SOX2 to mediate proliferation. Our data suggest that corneal stiffness-induced differentiation involves activation of SMAD2/3 and repression of YAP. The mechanism may involve the TGFβ ligand-mediated phosphorylation of SMAD2/3 that drives its nuclear localization and induces the transition to a differentiation regulatory network. TGFβ is known to bind a latency-associated peptide and the latent TGFβ binding-protein-1 (Shi *et al*, 2011) and αvβ6 integrin dependent stiffness induced contractility has been shown to liberate TGFβ in epithelial cells (Giacomini *et al*, 2012). Interestingly, αvβ6 is also expressed by corneal epithelial cells (Stepp, 2006) and they were also found to secrete latent-TGFβ- binding protein through extracellular vesicles (Han *et al*, 2017). A very interesting open question is whether LSCs regulate the stiffness of their niche? While few recent studies focused on the biomechanics of epithelial cells *in vivo* (Bhattacharya *et al*, 2022), the mechanobiology of stromal mesenchyme represents another frontier. The question of how do fibroblast, immune, and other niche cells sense and respond to mechanical strain is open. Interestingly, changes in ECM stiffness can modulate the morphology, cytoskeletal organization, and subcellular pattern of force generation in corneal stromal cells treated with TGF-β1 (Maruri *et al*, 2020).

The regulation of YAP may be complex and involve the integration of diverse signals. The activation of LATS1/2 by stiff matrix coincided with YAP phosphorylation and cytoplasmic localization. However, LSCs that were co-cultured with NIH-3T3-J2 feeder cells on a plastic dish (Giga Pascals) expressed high levels of nYAP in the colony periphery (Fig. 1B), suggesting that feeder cells provide an essential signal that controls YAP localization and stemness. Indeed, the induction of differentiation by calcium (on a plastic dish without feeder cells) was more forceful than differentiation induced by stiffer gel (compare Fig. 1D with Fig. 5A). Altogether, these observations align with a model where the biomechanical signals act in concert with other external signals mediated by niche factors.

In conclusion, this study suggests that biomechanical signals provide a critical cell fate determination signal to LSCs under homeostasis and regeneration. It will be of interest to investigate the relevance of these findings to corneal pathologies, such as LSC deficiency or keratoconus, that may involve changes in ECM rigidities (Fatima *et al*, 2007; Gouveia *et al*, 2019). While this study provides a potential explanation for the positive effect of ROCK and TGFβ inhibitors on the long-term expansion of epithelial cells (Zhang *et al*, 2018), further study of LSC mechanotransduction pathways is needed to improve LSC expansion, allowing their optimal application in regenerative medicine.

## Methods

### Animal handling

Animal care and use conformed to the ARVO Statement for the Use of Animals in Ophthalmic and Vision Research. The mouse strains K15-GFP (#005244), R26R-Confetti (#013731), K14-Cre^ERT^ (#005107) were from JAX (Bar Harbor, ME). Lox^OE^ was previously described (Gabay Yehezkely *et al*, 2020). Cre recombinase activity was induced by injecting intraperitoneally (200 μl), 4 mg/day of Tamoxifen (T5648, Sigma, St. Louis, MO) dissolved in corn oil for 3-4 consecutive days, as previously reported (Amitai-Lange *et al*, 2015). For wounding, mice were anesthetized (2% Isoflurane) and injected intramuscularly with analgesic Buprenorphine (0.03 mg/ml, 50 µl). Limbal and central corneal wounding (2 mm diameter) was performed using an ophthalmic rotating burr (Algerbrush) under fluorescent binocular. The wounded corneas were stained (1% fluorescein) and the wounded area was imaged and quantified (NIS-Elements analysis D and ImageJ software). For sub-conjunctival injection, the bulbar conjunctiva was pulled using forceps and 15 μl of Verteporfin (Sigma, ML0534) was injected under the binocular using a 30-gauge needle connected to 1 mL syringe. Slow injection into the space between the conjunctiva and the sclera was performed to create a ballooning effect in the peri-limbal conjunctival zone.

### Cell culture, differentiation, transfection, and Real-time PCR

Human limbal rings from cadaveric corneas were obtained under the approval of the local ethical committee and declaration of Helsinki from at least 3 corneas from 3 different donors. The epithelium was separated from the underlying stroma following incubation with dispase II (GIBCO, Life Technologies, USA). Cells were cultured at 37C, 5% CO2, and 20% O2. For clonogenicity assay cells were grown in co-culture with mitomycinized growth-arrested J2-NIH 3T3 cells in Green medium (60% DMEM (GIBCO), 30% DMEM

F12 (GIBCO), 10% FCII serum (Hyclone), 1mM L-Glutamine (Biological Industries), 1mM Sodium Pyruvate (Biological Industries), 0.2mM Adenine (Sigma), 5mg/ml Insulin (Sigma), 0.5mg/ml Hydrocortisone (Sigma), 10mM Choleratoxin (Sigma), 10ng/ml EGF (Peprotec)) and split in 80% confluence. For efficient transfection and controlled calcium-induced differentiation, cells were switched to a defined medium with supplements (SCMK001, Millipore, United States) containing 1% penicillin/streptomycin and low calcium (150µM). Cells were seeded and grown to 80-100% confluency for differentiation and then switched to high (1.2mM) calcium for up to 1 week. Cells were grown on defined media and collected at indicated time points after treatment with vehicle or indicated factors Blebbistatin (para-nitroblebbistatin, Optopharma Ltd), or Verteporfin (Sigma, ML0534) and taken for the Clonogenicity test.

MEFs and WI-38 cells were serum-starved for 15 hours and were kept in suspension in a serum-free medium for 30 minutes before seeding on fibronectin-coated PDMS gels to synchronize YAP localization to the cytoplasm. Cells were cultured in growth medium for 12 hours, fixed with 4% PFA, and then underwent immunostaining. Cells were imaged by ImageXpress® Micro Confocal (IXMC) microscope (Molecular Devices) with a 20X objective. Images were analyzed by the Translocation-Enhanced application module in MetaXpress® Software. In this module, an “inner region” shrunk in 0.5 µm from the detected nucleus was set to indicate the nuclear area, and an “outer region” expended out 0.5 µm from the detected nucleus was set to indicate the cytoplasmic area with a width of 2 µm. The Pearson’s correlation coefficient of the pixel intensity of YAP stain and nucleus stain in the two regions was calculated by the algorithm in the software. Correlation value of 1.0 indicates that the two stains overlap perfectly; –1.0 indicates complete lack of overlap between the two stains; and 0 indicates that the stains are independent. Cells with a correlation coefficient above 0.65 were considered YAP localized inside the nucleus. Nuclear-to-cytoplasmic (N/C) YAP average intensity was the average YAP intensity in inner regions divided by the average YAP intensity in outer regions.

For TUNEL, ApopTag® Red In Situ Apoptosis Detection Kit (S7165, Sigma) was used and staining were done according to the manufacturer’s instruction. For transfections, cells were seeded on plastic dishes and the next day transfected (Lipofectamine RNAimax, ThermoFischer) with 50nM esiRNA against EGFP (EHUEGFP, Sigma) or esiRNA against YAP (EHU113021, Sigma) or with following siRNA against YAP1 GACAUCUUCUGGUCAGAGA dTdT, YAP2 CUGGUCAGAGAUACUUCUU dTdT or ctl siRNA (Sigma, SIC001). Cells were collected 48-72 hrs after transfection. Real-time PCR analysis cells were washed with cold PBS and RNA was isolated using TRI-Reagent (Sigma) according to the manufacturers’ instructions. cDNA was prepared by reverse transcription polymerase chain reaction (RT-PCR) using the Qscript cDNA synthesis kit (Quantabio) according to the manufacturer’s instructions. Quantitative real-time polymerase chain reaction (qPCR) was performed with FastSYBR green master mix (Thermo). Samples were cycled using StepOnePlus (Applied Biosystems) qPCR system. Relative gene expression was normalized to GAPDH and calculated according to the DDCT method for qPCR.

### Western blots and immunostaining

Primary human limbal SCs were washed with cold phosphate-buffered saline (PBS) twice, and lysates were obtained in RIPA buffer (Tris-HCl 10 mM, 10 mg/ml deoxycholate, 1% NP40, 1% SDS, 150 mM NaCl, and protease inhibitors cocktail [Roche, Mannheim, Germany]). Total protein was subjected to polyacrylamide gel electrophoresis in the presence of sodium dodecyl sulfate. Proteins were separated on 12% polyacrylamide gel and transferred to nitrocellulose membranes (Bio-Rad) as reported 30-32. The membranes were blocked with trizma base buffer supplemented with 0.1% tween 20 (TBST, Sigma, USA) containing 5% milk (Bio-Rad, USA) and probed with the primary antibody in blocking solution at 4°C, overnight, followed by three washes with TBST. Furthermore, the membranes were exposed to peroxidase-conjugated goat anti-mouse IgG or peroxidase-conjugated goat anti-rabbit IgG or peroxidase-conjugated donkey anti-goat IgG (all at 1:3,000) for 1 hour at room temperature and washed three times with TBST. Protein bands were visualized with EZ-ECL Enhanced Chemiluminescence Detection Kit (Biological Industries, Israel).

For in-vitro staining, cultured epithelial cells or mouse fibroblasts (Wolfenson *et al*, 2016; Meacci *et al*, 2016) were grown on glass coverslips or silicone substrates and fixed in 4% paraformaldehyde (PFA) (Sigma) for 15 minutes and then permeabilized with 0.1% Triton X-100 (BioLab) in PBS for 10 minutes. Blocking was done with bovine serum albumin (Biological Industries), 3% donkey serum (Jackson) and 0.1% Triton X-100 for at least 30min. Following these treatments, cells were incubated for overnight with primary antibody at 4 degrees and further incubated with secondary antibodies (1:500) for 1 hour followed by 4′,6-diamidino-2-phenylindole (DAPI) or phalloidin 488 (1:400, Thermo Fisher, A12379) staining, and mounting (Thermo Scientific).

For wholemount staining, the cornea was isolated and fixed (2% formaldehyde) for 2 hours, room temperature followed by permeabilized (0.5% Triton, 5 hours). Blocking was done for 1 hour (0.1% TritonX-100, 2% Normal donkey serum, 2.5% BSA) and then incubated with primary antibody overnight, 4°C on a shaker and further incubated with secondary antibodies (1:500, 1 hour), followed by DAPI, tissue flattening under a dissecting binocular and mounting (Thermo Scientific). For EdU staining, a Single intraperitoneal injection of 200 μl (7.5 mg/ml) EdU (Invitrogen) was performed and 6 hours later, tissues were processed. Isolated corneas were fixed (2% PFA, 1 hour) and stained (Click-iT, Invitrogen) according to the manufacturer’s instructions followed by wholemount staining protocol (described above) for other markers staining.

Paraffin sections (5µm) of mouse cornea and human corneas were used and processed for immunohistochemistry. Paraffin sections were deparaffinized by heating for 1 hour at 60°C, then rehydrated twice in Xylene for 5 minutes, followed by two incubations in 100% Ethanol for 5 minutes. Suppression of endogenous peroxidase activity was achieved by incubation in Methanol with 1% H_2_O_2_, incubation in 70% Ethanol for 2 minutes, followed by a rinse in Distilled water. Antigen retrieval was performed with an unmasking solution (Vector Laboratories, H3300), followed by blocking (10% goat serum), and incubation with a primary antibody (overnight, 4C), secondary antibodies (Universal Immuno peroxidase Polymer anti-rabbit/mouse, a ready-made solution, 1 hour) followed by substrate addition (AEC solution), Hematoxylin staining and mounting (Thermo Scientific).

Images were taken by Nikon Eclipse NI-E upright microscope and Zeiss LSM880 confocal microscope. For quantification three to five different fields were imaged from different experiments and the indicated mean fluorescence intensity was calculated by ImageJ software.

### Antibody Table

| Antibodies | Source |
| --- | --- |
| YAP (1:200) | Santa cruz biotechnology, sc-101199 |
| YAP (1:200) | Abnova, H00010413-M01 |
| pYAP (1:1000) | Cell signaling, #13008 |
| SMAD2/3 (1:400) | Abcam, ab202445 |
| pLATS1 (1:400) | Enco, 50-198-8119 |
| K15 (1:1000) | Santa cruz biotechnology, sc-47697 |
| K12 (1:400) | Abcam, ab185627 |
| GAPDH (1:1000) | Cell signaling, #2118 |
| PHALLOIDIN (1:400) | Thermo Fisher, A12379 |
| VINCULIN (1:300) | Thermo Fisher. #700062 |
| TP63 (1:400) | Santa Cru, sc-8431 |
| K4 (1:100) | Abcam, ab183329 |
| CD63 (1:100) | Santa cruz biotechnology, sc-5275 |
| GPHA2 (1:1000) | Santa Cruz Cat, #sc-390194 |
| EDU | Invitrogen, E10187 |

### Tissue stiffness measurement by Atomic Force Microscopy (AFM)

Eyes were collected from 2 and 6 weeks old mice (C57BL/6J or Lox^OE^) following CO_2_ sacrifice. Corneas with surrounding limbus were isolated as previously described (Amitai-Lange *et al*, 2015), then treated for 30min in 2.5mM EDTA in PBS at 37C to allow epithelium removal. Next, samples were prepared by dissecting radially the tissue in two halves, each of them containing both the limbus and the cornea. The two halves were placed on SuperFrost Plus adhesion slides (Thermo Scientific) side by side and at 180 degrees to each other to have for both the areas of interest, the limbus and the cornea, free access to the sample for the AFM probe. The samples were allowed to adhere to the charged glass for 20’, then the fragment of glass containing the mounted tissues was cut out from the glass slide and glued on the bottom of a 50 mm dish (Willco Glass Bottom Dish). Before measurements the specimen was first rinsed and after covered with 4 ml of PBS 1x. After determining both the deflection sensitivity of the AFM system in PBS 1x using a clean glass slide fragment glued on Willco Glass Bottom Dish and the spring constant of the AFM cantilever by means of the thermal tune method, the sample was mounted on the AFM system and after thermal stabilization, for each limbus and the corresponding cornea (n=3), a minimum of 3 different areas were analyzed using the “Point and Shoot” method, collecting on average 100 force-distance curves at just as many discrete points spaced by at least 20 μm. Force-distance curves were collected on samples using a velocity of 2 μm/s, in relative trigger mode and by setting the trigger threshold to 1 nN. The apparent Young’s modulus was calculated using the NanoScope Analysis 1.80 software (Bruker Nano Surfaces, Santa Barbara, CA, USA) applying to the force curves, after the baseline correction, the Hertz spherical indentation model using a Poisson’s ratio of 0.5. All the force-distant curves having a not clear base line, a maximum above or below 1nN or a change of slope in the region of the fitting (minimum and maximum force fit boundary 0% and 25%, respectively) were rejected and not considered for the analysis. Only the apparent Young’s modulus values corresponding to a fit with R^2^> 0.85 were considered for the analysis. The tissue mechanical proprieties were obtained by using a Bioscope Catalyst (Bruker Nano Surfaces, Santa Barbara, CA, USA), coupled with an inverted optical microscope (Leica DMI6000B, Leica Microsystems Ltd., UK). The force-distance curves needed to calculate the apparent Young’s modulus were collected using a Borosilicate Glass spherical tip (5 μm of diameter) mounted on a cantilever with a nominal spring constant of 0.06 N/m (Novascan Technologies, Ames, IA USA).

### Silicone substrates preparation

To obtain silicone substrates of different stiffness, the elastomer CY52-276 (Dowsil) was used, with various ratios of silicone base and crosslinking agent (components A/B): 1:1.8 for 8 kilo Pascal (kPa), and 1:2.5 for 20 kPa. After thorough mixing of both components, air bubbles were eliminated by application of vacuum for 30 min. Gels were spread on glass-bottom dishes followed by incubating at 70 °C for 2 h. Then the gels were sterilized by 30 min immersion in 70% ethanol. The silicone gels’ surfaces were coated with 10 μg/ml fibronectin for at least 1 h before seeding cells.

### Pillar arrays fabrication

Pillar fabrication was done by pouring PDMS (Sylgard 184, Dow; mixing ratio – 10:1) into silicon molds (fabricated as previously described (Ghassemi *et al*, 2012) with holes at fixed depths and distances. The molds were then placed, face down, onto glass-bottom 35 mm dishes (#0 coverslip, Cellvis) which were incubated at 65°C for 12h to cure the PDMS. The molds were next peeled off while immersed in ethanol to prevent pillar collapse. The ethanol was then replaced by serial dilutions with PBS, and human plasma full-length fibronectin (Merck) was added to the dish at a final concentration of 10 µg/µl for a 1h incubation at 37°C. Next, residual fibronectin was washed away by replacing the buffer to defined media with 20 mM HEPES.

All pillars had a diameter of 2 µm, and heights of 5.3 or 13.2 µm. We used 2 μm diameter pillars as these can be used to measure the long-term time-dependent forces that are generated after initial formation and reinforcement of the adhesions (Feld *et al*, 2020). The center-to-center spacing between pillars was 4 μm. Pillar bending stiffness, k, was calculated by Euler–Bernoulli beam theory:

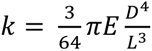

where D and L are the diameter and length of the pillar, respectively, and E is the Young’s modulus of the material (=2 MPa for the PDMS used here).

### Pillar displacement measurements

For measuring forces by K15-positive and K15-negative cells, the cells were grown in low and high-calcium media, respectively, for 4 days. On the day of the experiment, the cells were trypsinized, centrifuged with growth medium, and then resuspended and pre-incubated in defined media with HEPES at 37°C for 30 min before plating them on the 5.3 µm fibronectin-coated pillars.

Time-lapse imaging of cells spreading on the pillars was performed using an inverted microscope (Leica DMIRE2) at 37°C using a 63x 1.4 NA oil immersion objective. Brightfield images were recorded every 10 seconds with a Retiga EXi Fast 1394 CCD camera (QImaging). The microscope and camera were controlled by Micromanager software (Edelstein *et al*, 2010). In all cases, we imaged cells that were not in contact with neighboring cells when plated on the substrates. For each cell, a movie of 1-3 hours was recorded. To minimize photodamage to the cells, a 600 nm longpass filter was inserted into the illumination path.

Tracking pillar movements over time was performed with ImageJ (National Institutes of Health) using the Nanotracking plugin, as described previously (Ghassemi *et al*, 2012). In short, the cross-correlation between the pillar image in every frame of the movie and an image of the same pillar from the first frame of the movie was calculated, and the relative x- and y-position of the pillar in every frame of the movie was obtained. To consider only movements of pillar from their zero-position, we only analyzed pillars that at the start of the movie were not in contact with the cell and that during the movie the cell edge reached to them. Drift correction was performed using data from pillars far from any cell in each movie. For each pillar, the displacement curve was generated by Matlab (MathWorks).

### Statistical analysis

All experiments were performed at least three times. All quantifications represent the mean ± standard error of the mean (s.e.m.) or as indicated in the legends. Images are representative of experiments that have been repeated at least three times. Data were tested for normality where applicable by the Shapiro-Wilk method. Group comparison was performed using a two-tailed unpaired Student’s *t* test or Mann Whitney test as indicated in the legends. Multiple groups comparison was performed by ANOVA followed by Tukey’s test. Differences were considered to be statistically significant from a *p*-value below 0.05.

## Acknowledgment

We thank D. Aberdam for the critical reading of the manuscript. RSF has received funding from the Israel Science Foundation (1308/19 and 2830/20), Rappaport Family Institute for Research in Medical Sciences, NIH-exploratory R21 (800040), European Union’s Horizon 2020 research & innovation programme (828931). HW acknowledges support from the Israel Science Foundation (1738/17) and from the Rappaport Family Institute for Research in Medical Sciences; HW is an incumbent of the David and Inez Myers Career Advancement Chair in Life Sciences. CF was supported by INCA (Institut national du cancer) PL-Bio #2019-11/368/NI-HO, and the French Government (National Research Agency, ANR) through the “Investments for the Future” programs LABEX SIGNALIFE ANR-11-LABX-0028-01 and IDEX UCAJedi ANR-15-IDEX-01. The atomic force microscopy of IRCAN was supported by the Association pour la Recherche sur le Cancer (ARC), by the Infrastructures en Biologie Santé et Agronomie (IBiSA) and by the Conseil Départemental 06 de la Région Provence Alpes-Côte d’Azur. PH has received funding from the Israeli Science Foundation grant number 1111/18.

## Author Contribution

SB was involved in conceptual and experimental design, performed and interpreted the majority of the experiments, prepared the figures, analyzed data, and participated in the manuscript writing; AM performed the pillar experiments, analyzed data, and participated in ongoing discussions about the study; SP, CF performed AFM experiment, analyzed data, prepared the figures, and participated in the discussion and manuscript writing; SD, EC, AA, WN, SD, IM, OJ, AAL performed experiments and provided data; MM, SS performed subconjunctival injections; AK, PH provided materials and participated in discussions and manuscript writings; HW and RSF were involved in conceptual and experimental design, data interpretation, and participated in manuscript writing. All authors approved the manuscript.

## Figure legends

**Figure S1.**
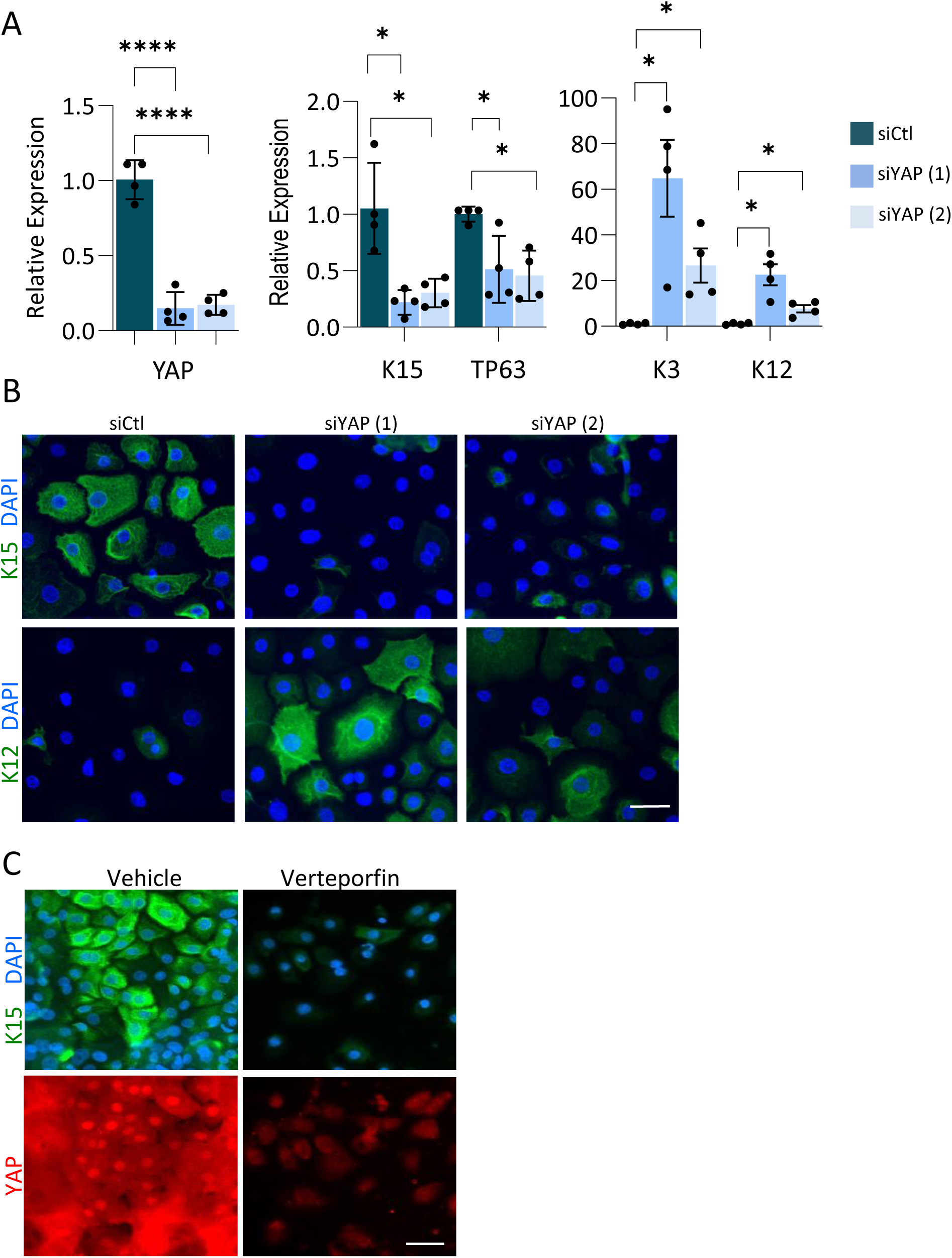
Knockdown of YAP by silencing RNAs or verteporfin induces LSC differentiation. (A-C) Primary human LSCs were grown on plastic dishes and transfected with two distinct silencing RNA sequences against YAP (siYAP) or with control sequence (siCtl). (A) The expression of the indicated genes was examined by real time PCR analysis, or by immunofluorescent staining (B) 48-72 hours post transfection. (C) Primary human LSCs were grown for 4 days on silicone substrate that mimics the rigidity of the limbus coated with fibronectin in the presence Verteporfin of Vehicle (control) and stained for K15 or YAP. Real-time PCR data was normalized to the housekeeping gene and is presented (mean ± standard error of mean, n=4 biological replicates) as fold increase compared to control sample and statistical analysis was performed by t-test (*, p < .05; **, p < .01; ***, p < .001, ****, p < .0001). Nuclei were detected by DAPI counterstaining. Scale bars are 50µm.

**Figure S2.**
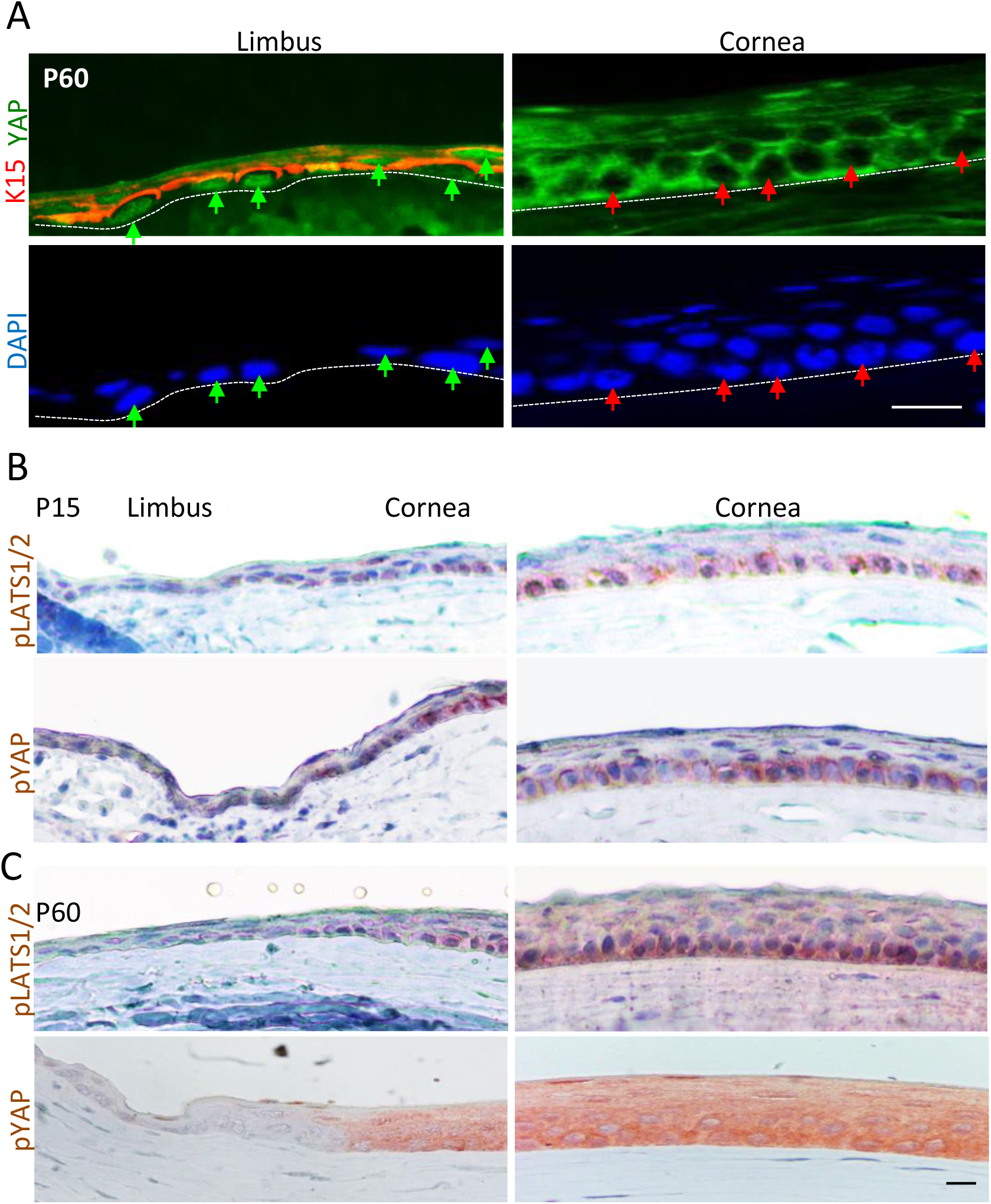
Analysis of YAP and Hippo pathway kinases. (A) Co-immunofluorescent staining of YAP and K15 in tissue sections of P60 murine cornea. Green arrows indicate nYAP in the limbus and red arrows indicate cYAP in the cornea compartment. (B-C) Immunohistochemistry of pYAP and pLATS1/2 was performed in the tissue sections of P15 (B) and P60 (C) of wild type murine cornea. Data represents 3 biological replicates. Scale bars are 50µm.

**Figure S3.**
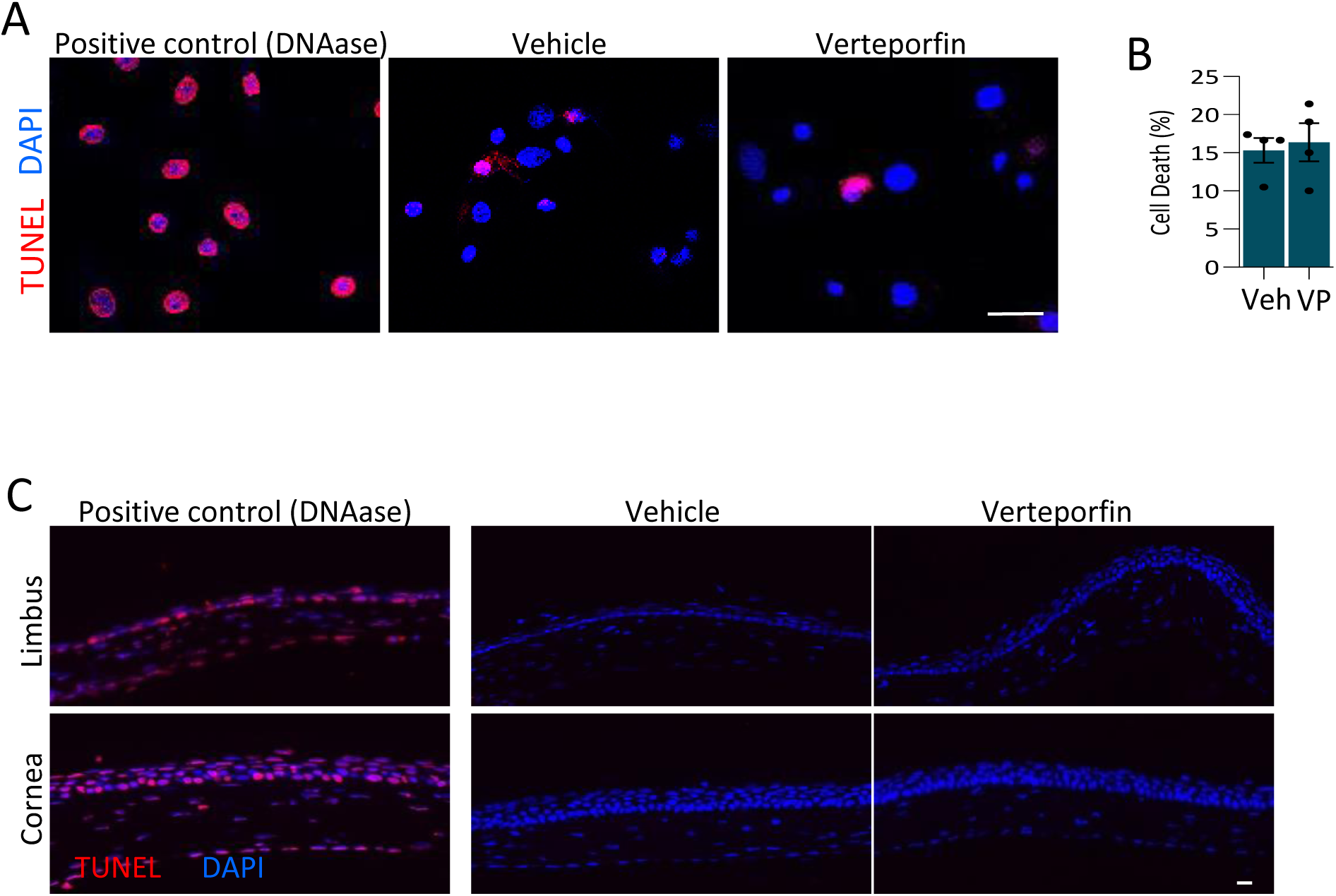
Verteporfin did not elicit cell toxicity. Primary LSCs were grown on silicone gels that mimic limbal rigidity in the presence of Verteporfin (VP) or Vehicle control. On the next day, cell death was assessed by TUNEL assay and cells were imaged (A) and quantified (B). (C) A daily sub-conjunctival injection (20µl) of Verteporfin (20µM) or vehicle (control) was performed for 4 days on K15-GFP transgenic mice. Tissue morphology was pictured following paraffin section preparation and apoptotic cells were detected by TUNEL assay. DNAase treatment served as positive control and nuclei were detected by DAPI counterstaining. Data represents 3 biological replicates. Scale bars are 50µm.

**Figure S4.**
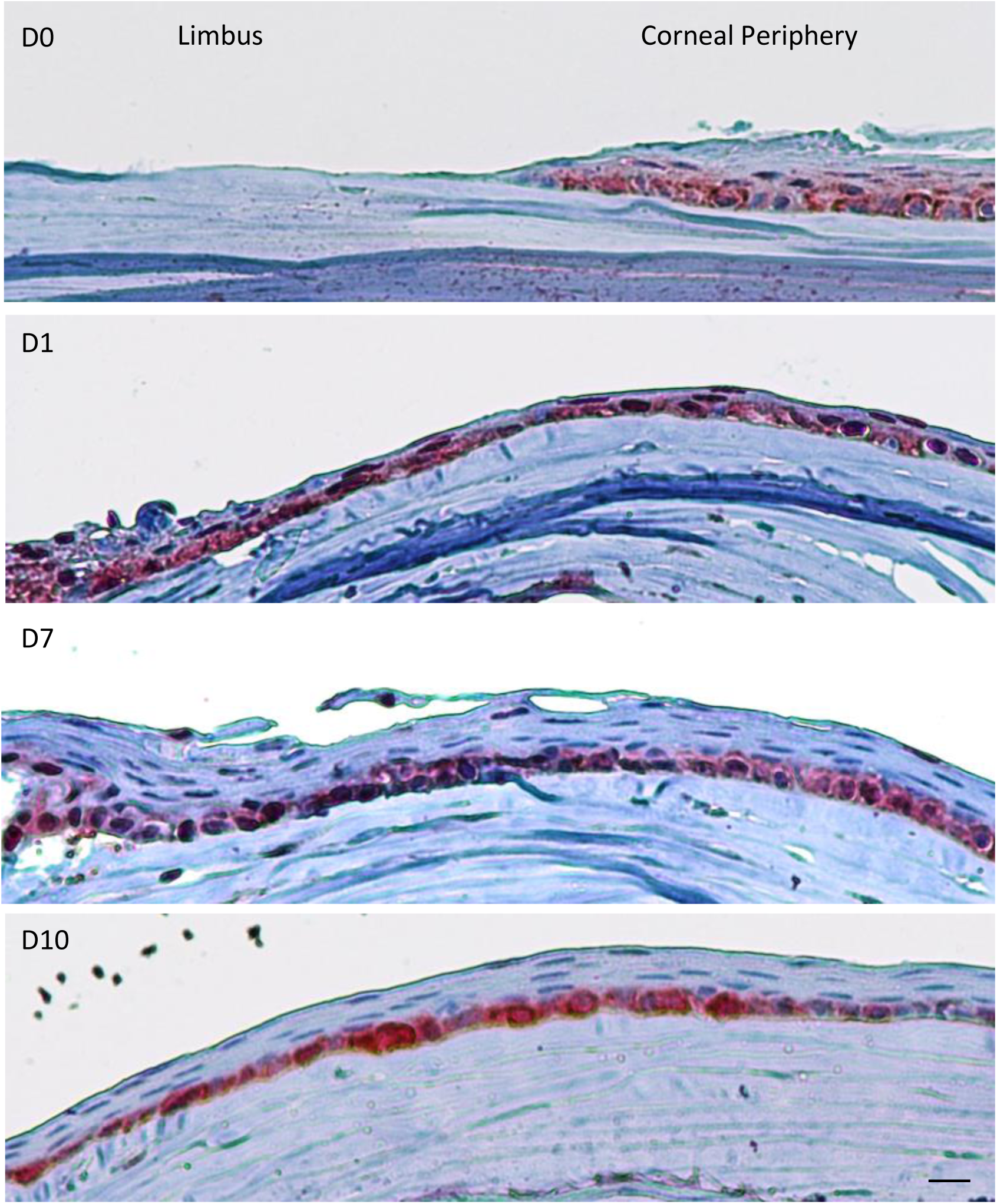
Analysis of YAP expression following limbal epithelial depletion. The limbal epithelium of adult mice was surgically removed (see Fig.3), mice were sacrificed in the indicated days post injury and immunohistochemistry was performed to detect YAP on paraffin limbus/cornea tissue sections. Data represents 3 biological replicates. Scale bars is 50µm.

**Figure S5.**
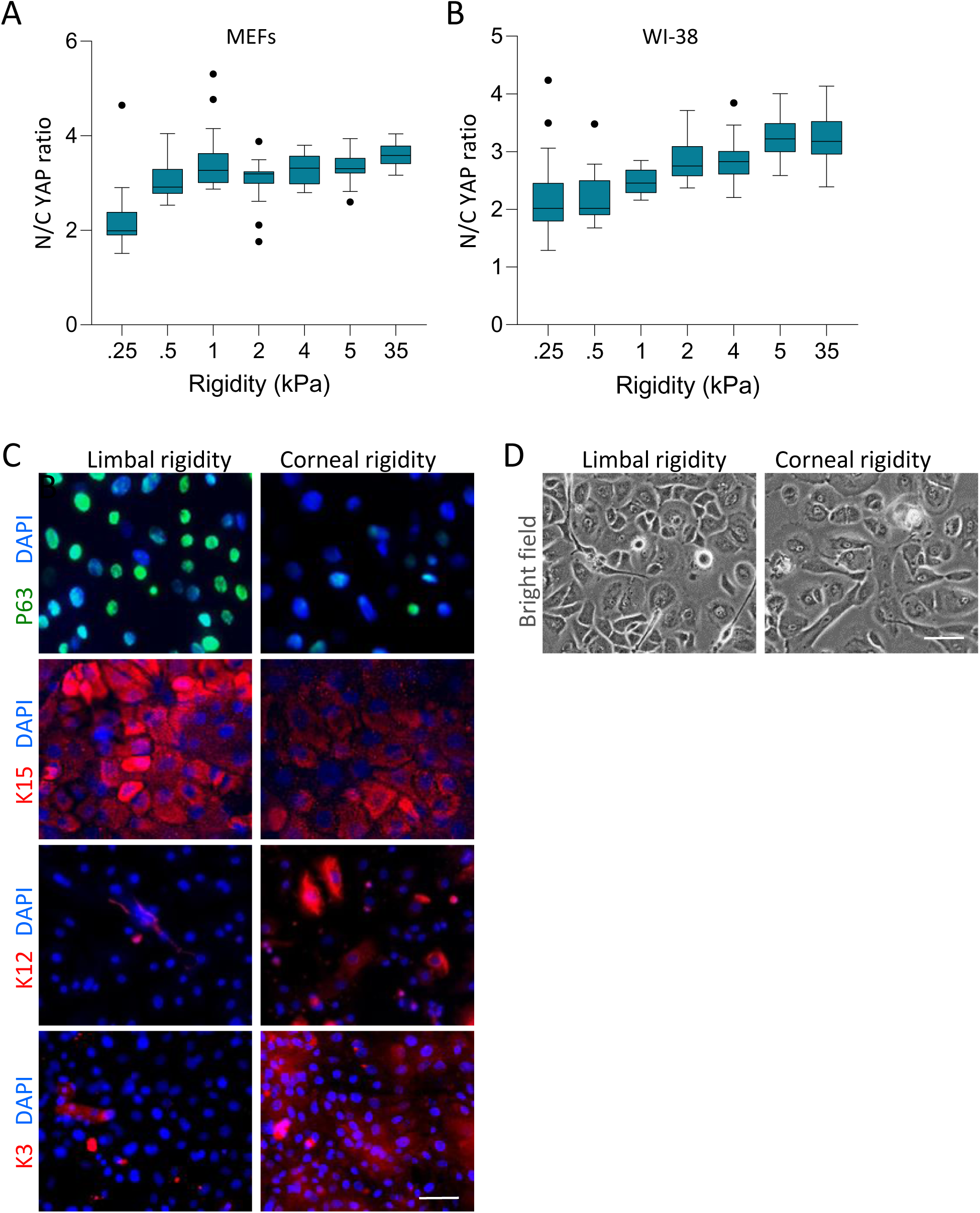
Corneal rigidity induces differentiation of human LSC. (A-B) Mean nuclear-to-cytoplasmic YAP intensity ratio of MEF (A) and WI-38 (B) cells on different rigidities for 12 hours. (C-D) Primary human LSCs were grown for 4 days on a silicone substrate that mimics the rigidity of the limbus (LR, 8kPa) or cornea (CR, 20kPa) coated with fibronectin. Cell differentiation state was examined by immunofluorescent staining of the indicated markers (C and brightfield images were taken (D). Data represents 3 biological replicates. Scale bars is 50µm.

**Figure S6.**
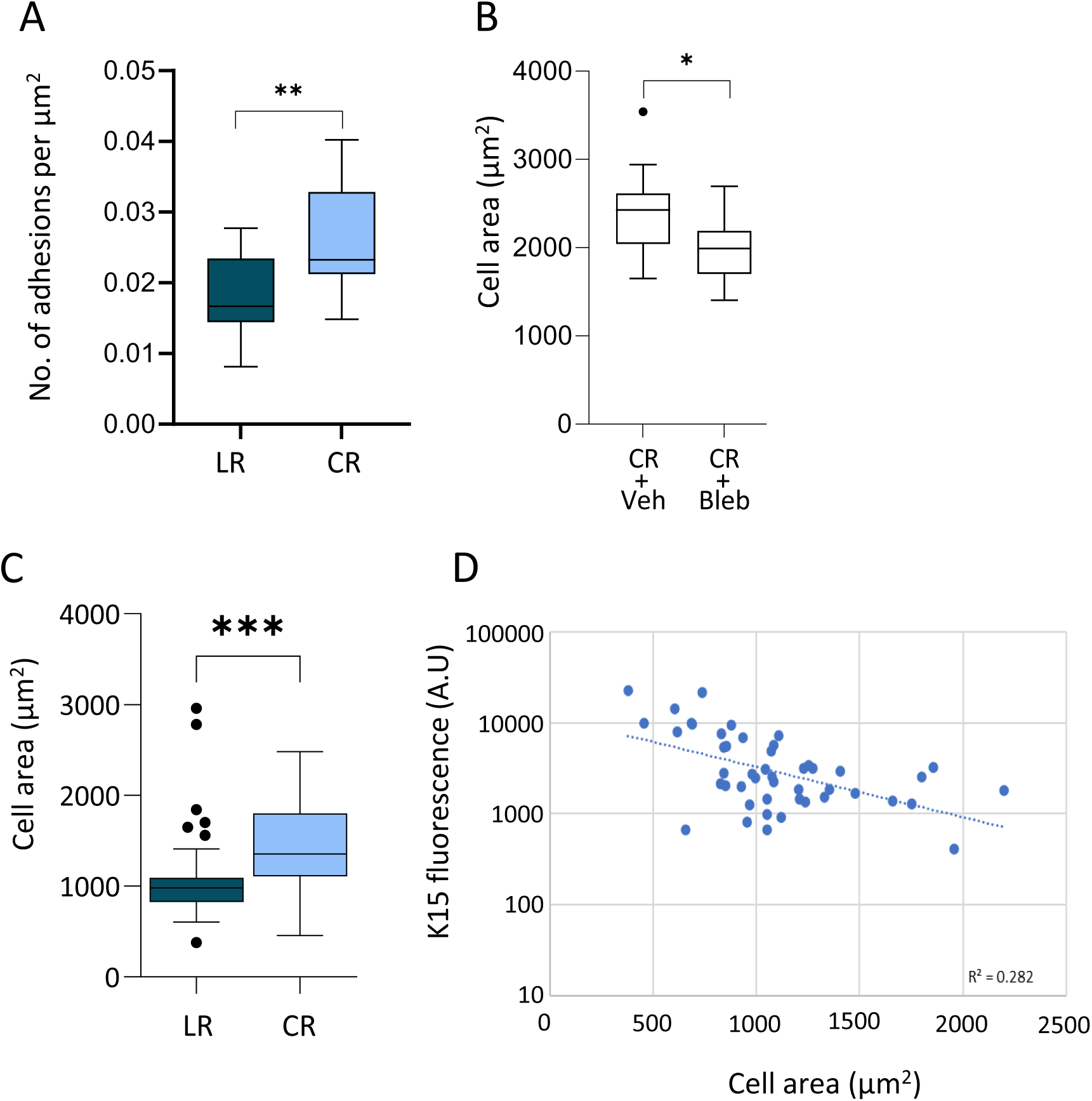
Analysis of cell area and adhesions on gels and pillars. (A) Analysis of the number of mature vinculin adhesions per µm^2^ in human LSCs that were grown on silicone gel with limbal rigidity (LR, 8kPa) or corneal rigidity (CR, 20kPa). (B) Analysis of the areas covered by LSCs on silicone gel with the indicated rigidity in the presence of Blebbistatin (Bleb), or vehicle (Control). (C) Areas of cells as measured using the phalloidin signal after overnight incubation on LR and CR pillars. (D) K15 immunostaining fluorescence signal as a function of cell area in cells plated overnight on the soft pillars. Data represents 3 biological replicates. The cell area is shown by the Tukey box-and-whisker plot followed by t-test (*, *p* < .05; **, *p* < .01; ***, *p* < .001, ****, *p* < .0001).

**Figure S7.**
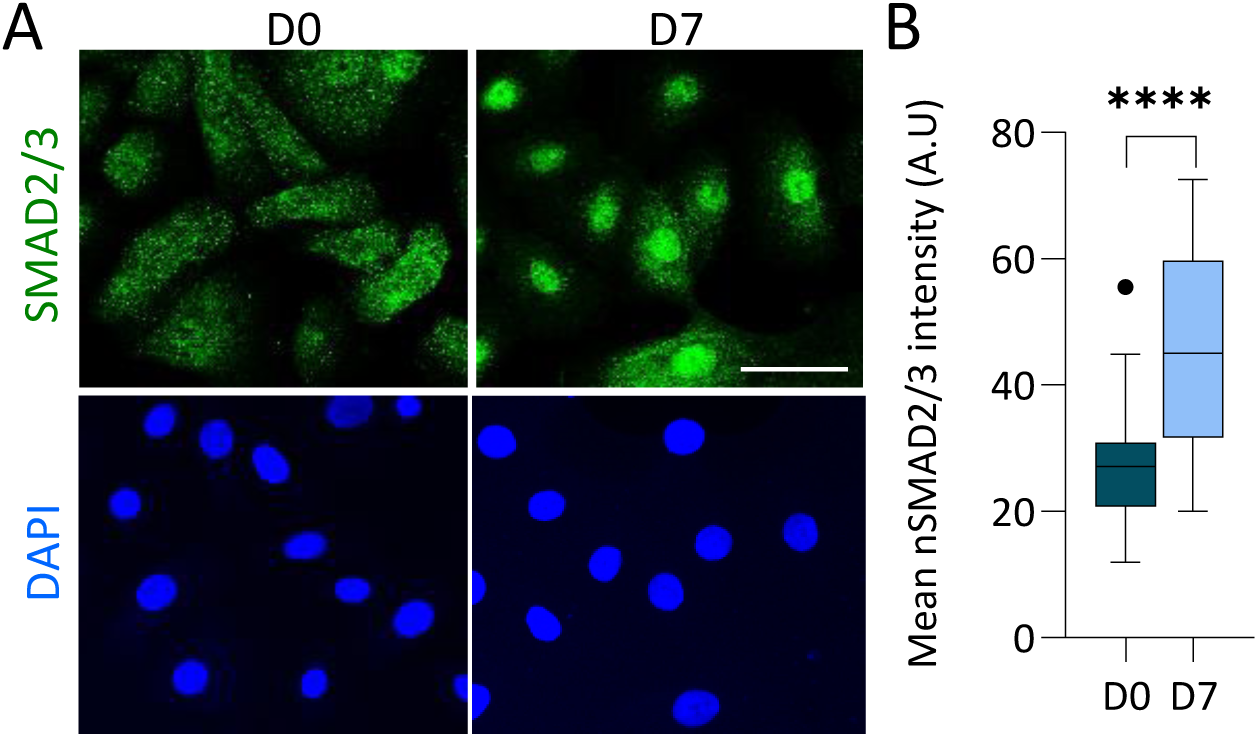
Nuclear SMAD2/3 is associated with differentiation. (A) Primary human LSCs were cultured on plastic culture dishes and maintained at low calcium (Day 0) or induced to differentiate in high calcium (Day7) and immunostained for SMAD2/3 (A) and mean nuclear SMAD2/3 intensity was quantified (n=3 biological replicates). (B) Mean nuclear SMAD2/3 intensity is shown by the Tukey box-and-whisker plot followed by t-test with Welch’s correction (*, *p* < .05; **, *p* < .01; ***, *p* < .001, ****, *p* < .0001). Data represents 3 biological replicates. Nuclei were detected by DAPI counterstaining. Scale bars is 50µm.

